# Structural basis for lipid and copper regulation of the ABC transporter MsbA

**DOI:** 10.1101/2022.08.04.502837

**Authors:** Jixing Lyu, Chang Liu, Tianqi Zhang, Samantha Schrecke, Nicklaus P. Elam, Georg Hochberg, David Russell, Minglei Zhao, Arthur Laganowsky

## Abstract

A critical step in Lipopolysaccharide (LPS) biogenesis involves flipping lipooligosaccharide, an LPS precursor, from the cytoplasmic to the periplasmic leaflet of the inner membrane, an operation carried out by the ATP-binding cassette transporter MsbA. Although MsbA has been extensively studied, the selectivity of MsbA-lipid interactions remains poorly understood. Here we use native mass spectrometry (MS) to characterize MsbA-lipid interactions and guide structural studies. We show the transporter co-purifies with copper(II) and metal binding modulates protein-lipid interactions. A 2.15 Å resolution structure of an N-terminal region of MsbA in complex with copper(II) is presented, revealing a structure reminiscent of the GHK peptide, a high-affinity copper(II) chelator. Our results demonstrate conformation-dependent lipid binding affinities, particularly for the LPS-precursor, 3-deoxy-D-*manno*-oct-2-ulosonic acid (Kdo)_2_-lipid A (KLA). We report a 3.6 Å-resolution structure of MsbA trapped in an open, outward-facing conformation with adenosine 5’-diphosphate and vanadate, revealing an unprecedented KLA binding site, wherein the lipid forms extensive interactions with the transporter. Additional studies provide evidence that the exterior KLA binding site is conserved and a positive allosteric modulator of ATPase activity, serving as a feedforward activation mechanism to couple transporter activity with LPS biosynthesis.

## Introduction

A defining feature of most Gram-negative bacteria is the presence of lipopolysaccharide (LPS) in the outer leaflet of the outer membrane.^1-3^ LPS contributes to the formation of an impermeable barrier that helps bacteria resist antibiotics and environmental stresses.^2^ Biogenesis of LPS commences in the cytoplasm with the production of the LPS-precursor lipooligosaccharide (LOS) followed by an orchestrated transport to the cell surface along with further modifications (**Fig 1A**).^4^ LOS contains a conserved lipid A moiety, a bisphosphorylated disaccharide of glucosamine (GlcN) with four to seven acyl chains, that is decorated with 3-deoxy-D-*manno*-oct-2-ulosonic acid (Kdo) sugar.^1^ Further decoration includes the attachment of a core oligosaccharide, which varies in different bacteria.^2^ Cytoplasmic LOS is flipped from the inner to the periplasmic leaflet of the inner membrane, an essential step carried out by the **A**TP-**B**inding **C**assette (ABC) transporter MsbA. As inhibition or deletion of MsbA is lethal,^5^the transporter has emerged as an attractive target for developing antibiotics. Small molecule MsbA inhibitors have recently been developed that vary in mode of action, such as trapping the transporter in an inward-facing (IF) conformation or mimicking substrate binding^.6-10^

**Figure 1.**
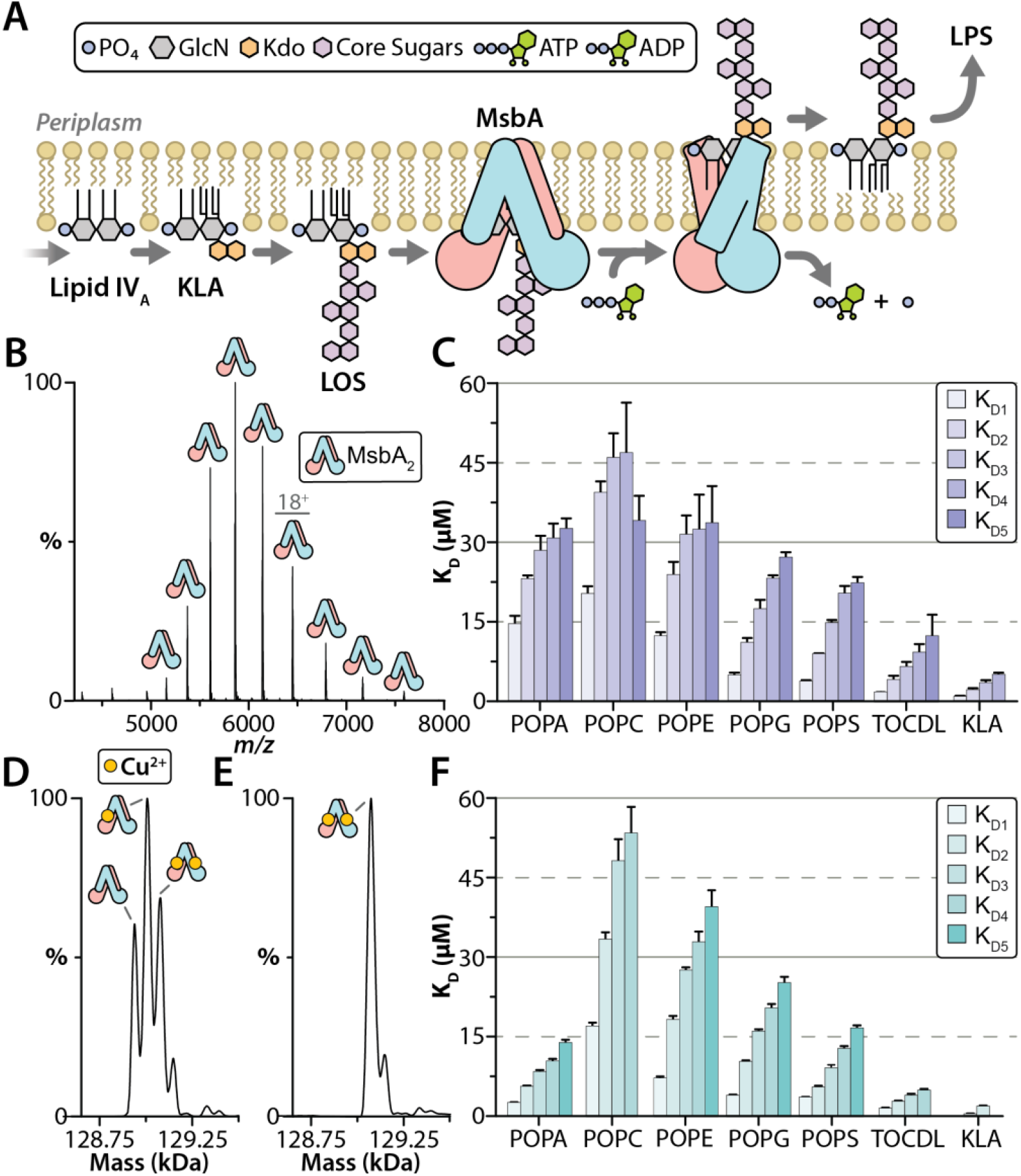
Copper(II) binding to MsbA modulates lipid binding affinity. A) Lipopolysaccharide biosynthesis commences in the cytoplasm to generate lipooligosaccharide (LOS). LOS is composed of conserved lipid A structure (gray), a bisphosphorylated disaccharide of glucosamine, modified with 3-deoxy-D-*manno*-oct-2-ulosonic acid (Kdo) sugar (orange) and core oligosaccharide (purple), of which is dependent on the bacteria. MsbA, powered by the hydrolysis of ATP, flips cytoplasmic LOS from inner to outer leaflet of the inner membrane, an essential step in LPS biogenesis. The flipped LOS is transported to the outer membrane along with additional modifications to become LPS. B) Native mass spectrum of optimized MsbA samples in C_10_E_5_ yields a well-resolved mass spectrum. C) Equilibrium dissociation constants (K_D_) for individual lipid binding events to partially loaded MsbA. D) Deconvolution of the mass spectrum shown in panel b. The different molecular species correspond to dimeric MsbA and different numbers of bound copper ions. E) Measured mass of MsbA after loading with copper(II) shows saturation of two binding sites. F) K_D_s for individual lipid binding events to MsbA fully loaded with copper(II). Reported are the mean and standard deviation (*n* = 3).

A number of studies employing an arsenal of biophysical techniques have provided mechanistic and structural insight into MsbA^.6,11-20^ MsbA forms a homodimer, each subunit consists of a transmembrane domain (TMD, six transmembrane helices per subunit) and a cytosol-exposed nucleotide-binding domain (NBD).^21^ The proposed mechanism by which MsbA translocates LOS across the bilayer involves several steps^.6,17^ Apo or adenosine 5’-diphosphate (ADP) bound MsbA populates an IF conformation with the NBDs separated in space, promoting entry and binding of the bulky LOS. The binding of LOS in the central, interior cavity and adenosine 5’-triphosphate (ATP) to MsbA promotes dimerization of the NBDs. ATP hydrolysis powers a conformational change to an outward-facing (OF) conformation, transporting LOS to the periplasmic side of the inner membrane. LOS and inorganic phosphate are released and MsbA cycles back to an IF conformation. Like many other ABC transporters, the ATPase activity of MsbA is stimulated in the presence of different substrates, particularly hexaacylated lipid A species, such as 3-deoxy-D-*manno*-oct-2-ulosonic acid (Kdo)_2_-lipid A (KLA).^22-24^ However, the structural basis for stimulation of MsbA ATPase activity by these substrates is poorly understood.

Native mass spectrometry (MS) has emerged as an indispensable biophysical technique to study membrane protein complexes and their interactions with lipids and other molecules.^25^ With the ability to preserve non-covalent interactions and native-like structure of membrane proteins in the gas phase^,26,^27 the technique has provided insightful information on various biochemical interactions including nucleotide, drug, peptide and lipid binding as well as yielding thermodynamic data for protein-protein, protein-ligand and protein-lipid interactions.^28-35^ In this study, we set out to characterize MsbA-lipid interactions in different conformational states. Native MS studies reveal MsbA co-purifies with copper(II) but also MsbA-lipid interactions are directly influenced by metal binding and protein conformation. Structural studies reveal a KLA binding site that has not been previously observed in other ABC structures. Our findings bring forth new insights into metal and lipid regulation of MsbA.

## Results

### Discovery of copper(II)-bound MsbA

As MsbA has been reported to co-purify with LPS and other lipids^,17,^36 our first objective was to optimize the purification of MsbA from *E. coli* for native MS studies. The mass spectrum of MsbA purified in the detergent n-dodecyl-β-D-maltopyranoside (DDM) was a broad hump (**Fig S1**), indicating a highly heterogenous sample, corresponding to a battery of co-purified small molecule contaminants. The sharp mass spectral peaks decorating the underlying hump, centered around 4500 *m/z*, correspond to monomeric MsbA, resulting from dissociation of the homodimer under the high activation, non-native conditions. After employing an established detergent screening method to optimize protein purification (**see Supplementary Text**),^37^ MsbA samples solubilized in the pentaethylene glycol monodecyl ether (C_10_E_5_) detergent exhibited a well-resolved mass spectrum (**Fig 1B**). Interestingly, different molecular species are measured, corresponding to dimeric MsbA and the addition of one to three ∼65 Da adducts (**Fig 1D and Table S1**). Analysis of MsbA samples using inductively coupled plasma mass spectrometry (ICP-MS) identified the bound adducts as copper (**Table S2-S3**). The addition of copper(II) to MsbA saturated the two binding sites (**Fig 1E**). Removal of excess copper(II) nor the addition of the copper(II) chelator, trientine^38^ reduced the amount of metal bound to MsbA (**Fig S2**). These results reveal MsbA has one high-affinity copper(II) binding site per subunit.

### Determination of MsbA-lipid binding affinities

To better understand MsbA-lipid interactions, we determined equilibrium binding constants for MsbA binding to different lipids. For these studies, we selected 1,1′,2,2′-tetraoleoyl-cardiolipin (TOCDL, 72:4) or phosphatidic acid (PA), phosphatidylcholine (PC), phosphatidylethanolamine (PE), phosphatidylglycerol (PG), and phosphatidylserine (PS) containing the acyl chain composition, 1-palmitoyl-2-oleoyl (PO, 16:0-18:1). We also included 3-deoxy-D-*manno*-oct-2-ulosonic acid (Kdo)_2_-lipid A (KLA), an LPS precursor known to stimulate MsbA ATPase activity.^22-24^ Except for PC, these lipids are found in *E. coli*. ^39^ MsbA partially and fully loaded with copper(II) was titrated with each lipid followed by recording their native mass spectra (**Fig S3-S6**). The mole fractions of apo and lipid bound states of MsbA were extracted from the deconvoluted MS data and used to determine the equilibrium dissociation constant (K_D*N*_) for the *N*^*th*^ binding event (**Table S4-S5**). For nearly all the lipids, MsbA loaded with copper(II) resulted in an enhancement in binding affinities, especially for POPA (**Fig 1C-f and S7**). The two tightest binding lipids for MsbA loaded with copper(II) were TOCDL (K_D1_ = 1.6 µM) and KLA (K_D1_ = 0.6 µM). POPG binding affinities were largely independent of the copper(II) bound state of MsbA. In short, these results demonstrate that MsbA not only binds selectively to lipids but is also dependent on the degree of copper(II) bound to the transporter.

### X-ray structure of the N-terminus of MsbA coordinated to copper

Two putative metal binding sites for a MsbA have been previously reported.^40^ Mutation of one of the putative sites (H562A and H576A) did not abolish copper(II) binding (**Fig S2**). After carefully inspecting MsbA structures with focus on histidine and cysteine residues, both of which are known to preferentially coordinate copper,^41^ we noted that in all MsbA structures the N-terminal histidine is not observed. Removal of the first four residues of MsbA abolished binding of copper(II) (**Fig 2A**), pinning down the metal binding site to the N-terminus. Additional studies show the truncated, copper(II)-free protein does not display altered ATPase activity (**Fig S8**). Interestingly, MD simulations^42^ show the N-terminus of MsbA in different structures are located near the inner membrane and in a region where LOS could pass before entering the interior cavity (**Fig 2B and S9**).

**Figure 2.**
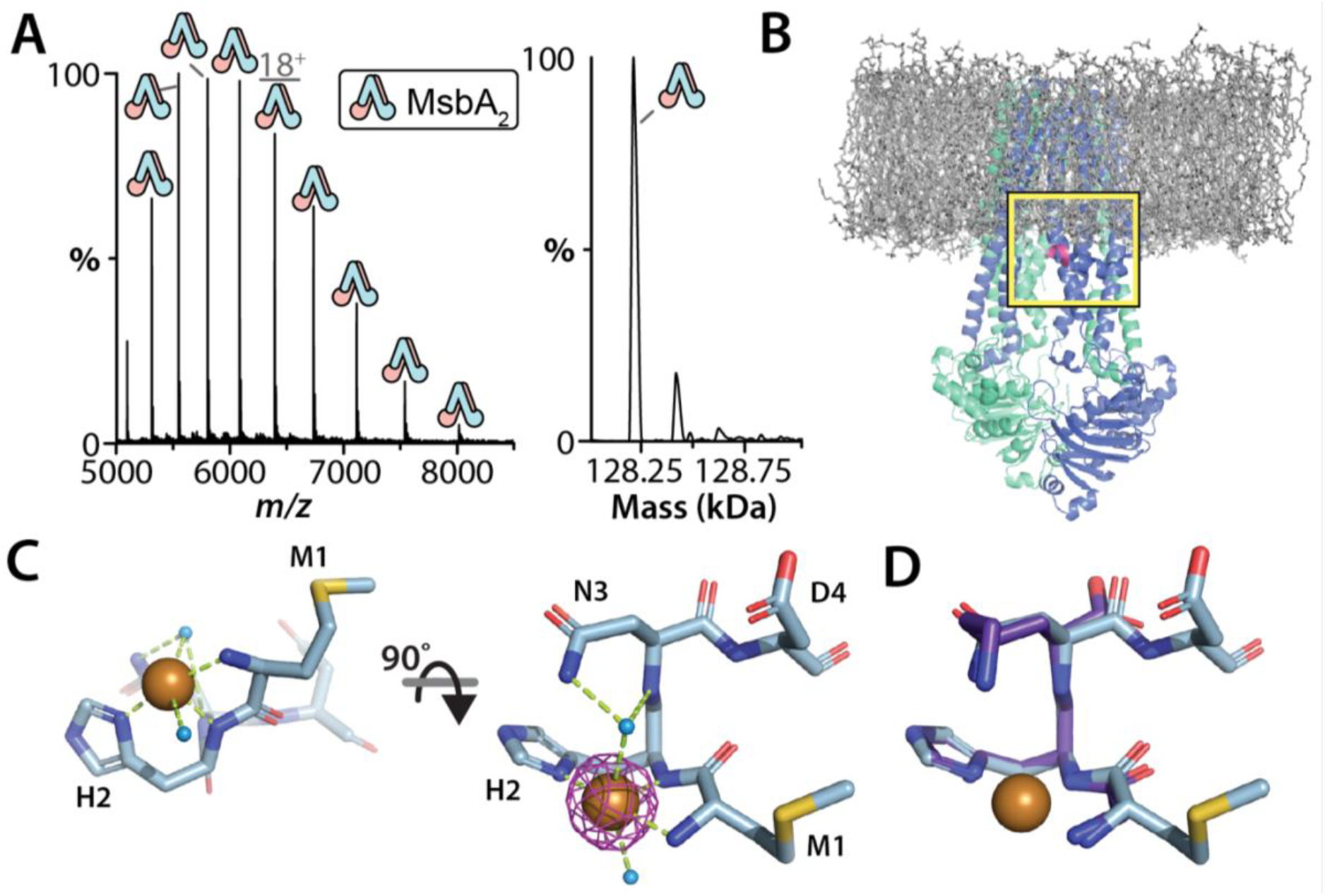
The N-terminus of MsbA binds copper(II) and its crystal structure. A) Native mass spectrum and deconvolution of MsbA with deletion of four N-terminal residues. No copper(II) is bound to the truncated transporter. B) Snapshot from a molecular dynamics simulation of MsbA in a 16:0 PC (DPPC) bilayer (PDB 6BPP downloaded from MemProtMD^42^). DPPC is shown in stick representation (grey). Protein is shown in cartoon representation with residues 4-8 colored pink. Yellow box highlights the location of the N-terminus relative to the inner membrane. C) Structure of the N-terminus of MsbA (residues 1-4) fused to the green fluorescent protein (GFP) coordinating copper(II). The N-terminal peptide is shown in stick representation with water (blue) and copper(II) shown as spheres. Bonds are shown as dashed lines (limon). Anomalous difference peak shown in magenta and contoured at 15 sigma. The structure of GFP is omitted for clarity. D) Alignment of the N-terminal MsbA peptide bound to copper(II) with GHK-copper(II) complex (CCDC-809108, C*α* colored purple).

To facilitate structure determination, the N-terminal sequence of MsbA was grafted onto proteins known to readily crystallize. Native MS shows these fusion proteins containing a fragment of the N-terminus of MsbA of variable length bound copper(II) and monomeric (**Fig S10**). One of the green fluorescent protein (GFP) fusions, containing residues 1-4 of MsbA, produced X-ray-grade crystals that led to structure determination at 2.15 Å resolution (**Table S6**). Resolved electron density for the N-terminus is observed along with strong anomalous signal for the bound copper ion (**Fig 2C and S11**) and adopts a structure of similar to that for the copper(II) coordinated GHK peptide, a naturally occurring high-affinity copper(II) chelator found in the blood plasma (**Fig 2D**).^43^ The copper(II) adopts a pseudo-octahedral coordination with the planar ligands comprised of the amine of M1, amide and sidechain of H2. An additional interaction is formed by the sidechain of D197’ from a symmetry related molecule (**Fig S10C**) and similar to the GHK-copper(II) structure, wherein is a C-terminal carboxylate. The axial ligands of copper(II) are water, one of which forms a bridge between copper and the sidechain and amide of N3. This coordination differs from the GHK peptide in the axial waters and N3 participating in a water bridge to the metal ion (**Fig 2D**).^44^ The difference between the two structures suggests the third position can be variable. Given the proximity of the N-terminus to the inner leaflet (**Fig 2B and S11**), the copper(II) bound structure could engage lipid headgroup, resulting in enhanced lipid binding affinity. Analysis of ABC transporter sequences, reveals more than 400 proteins contain a histidine in the second position, including some with an N-terminal sequence of MHK, and may have relevance for other ABC transporters.

### Characterizing lipid binding to vanadate-trapped MsbA

To determine the impact of MsbA conformation on lipid binding affinity, analogous experiments were performed using MsbA trapped in an open, OF conformation with adenosine diphosphate (ADP) and vanadate (VO_4_). The native mass measurement show that each subunit of the transporter is bound to copper(II), ADP, and VO_4_ molecules (**Fig 3A and Table S1**). The dissociation constants revealed that MsbA in the open, OF conformation displayed higher lipid binding affinity for a subset of lipids (**Fig 3D and S12-S14, and Table S7**). For example, in the presence of 0.4 μM of KLA, vanadate-trapped MsbA binds up to two lipids whereas the non-trapped protein binds only one KLA (**Fig 3B-C**). In particular, the binding affinity for KLA (K_D1_ = 0.3 µM) significantly increased by two-fold compared to the non-trapped protein. Interestingly, the change in K_D_ for each subsequent lipid binding event was significantly reduced, an indication of strong positive cooperativity. The binding of POPC and POPE are reminiscent of that for MsbA partially loaded with copper(II). POPA and POPG displayed an overall modest increase in binding affinity. Together, these results demonstrate that different conformational states of MsbA bind lipids with different affinities.

**Figure 3.**
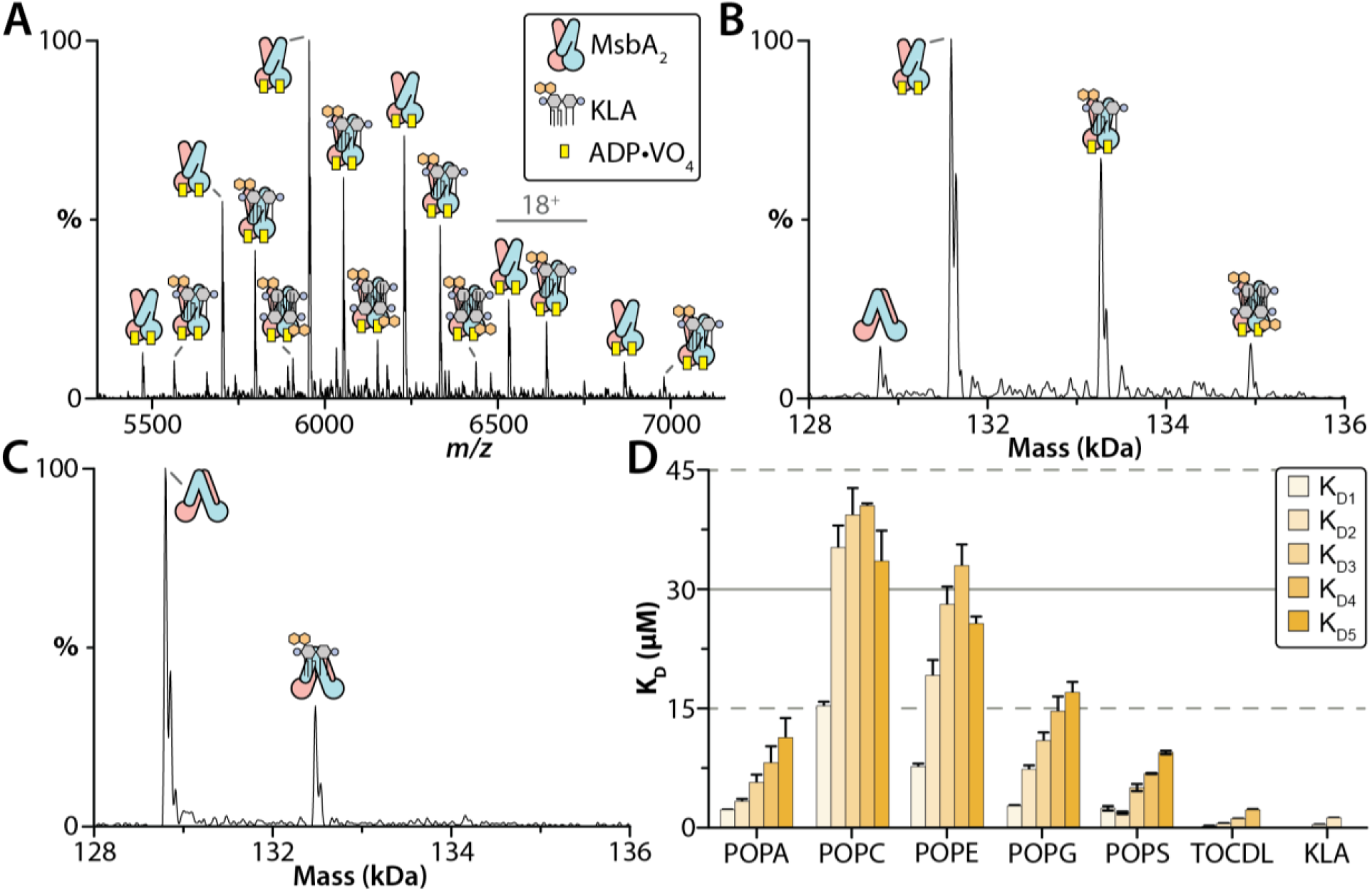
Biophysical characterization of lipid binding to MsbA trapped with ADP and vanadate. A) Representative mass spectrum of MsbA trapped with ADP·VO_4_ and in the presence of 0.4 μM KLA. B) Deconvolution of the mass spectrum shown in panel b. C) Deconvoluted mass spectrum of non-trapped MsbA in the presence of the same amount of KLA. A significant reduction in KLA binding to MsbA is observed. D) K_D_ values for individual lipid binding events to MsbA trapped with ADP and vanadate. Reported are the mean and standard deviation (*n* = 3).

### CryoEM structure of vanadate-trapped MsbA in complex with KLA

As KLA binds copper(II)-bound MsbA with an affinity greater than the other lipids, we prepared MsbA trapped with ADP and vanadate in the presence of 2 equivalents of KLA in C_10_E_5_ for cryo-electron microscopy (cryoEM) studies. The structure of the complex was determined to a resolution of 3.6 Å (**Fig 4A and S15, and Table S8**). The structure is similar to that of the vanadate-trapped MsbA from *S. typhimurium* but differs from the previously reported vanadate-trapped, occluded MsbA structures^,17,^45 largely in the TMD (**Fig S16**). Additional density is observed centered on TM5 in a region within the cytoplasmic leaflet of the inner membrane, in which KLA could be directly modeled into this density (**Fig 4A**). The location of KLA is different from the interior binding site^6,^17 and the putative, exterior binding site^19^ located on the opposite side of the inner membrane (**Fig S17**). The electrostatic surface of MsbA shows that the headgroup of KLA is bound to a large, basic patch (**Fig 4B**). Acyl chains of KLA could be modeled, which interact with the hydrophobic surface of MsbA (**Fig 4C**). Extensive interactions are formed between KLA and MsbA with an interface area of 2,779 Å^2^. The characteristic phosphoglucosamine (P-GlcN) substituents of LOS are coordinated by R238 on one side, and R188 and K243 on the other side (**Fig 4D**). In addition, R236, Q240, and K243 interact with one of the Kdo groups of LOS (**Fig 4D**). In addition, there is clear density for ADP·VO_4_ in the NBDs, coordinated by a network of conserved residues (**Fig 4E and S18**). In short, the new KLA binding site identified here may have role in regulating MsbA function.

**Figure 4.**
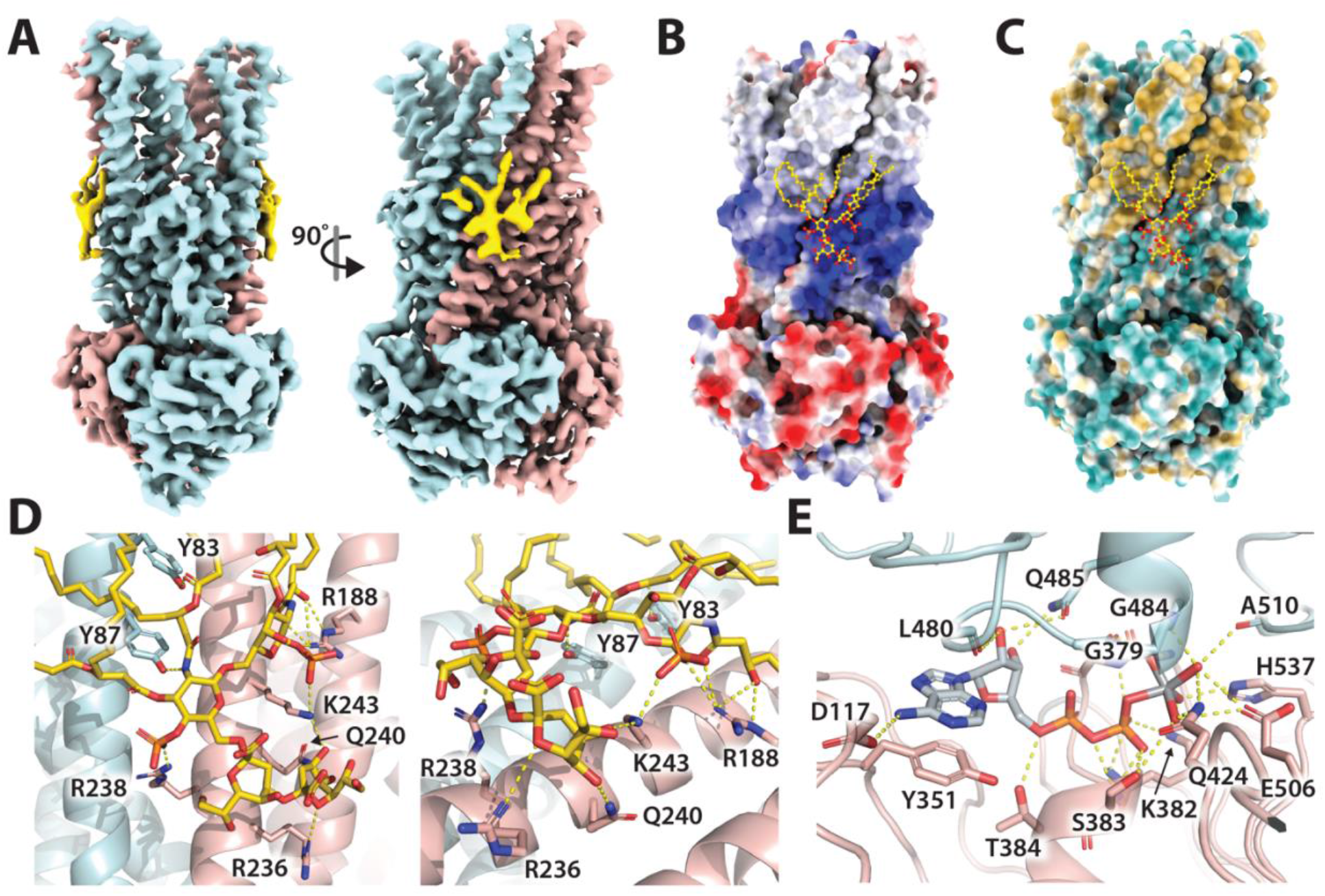
Structure of vanadate-trapped MsbA reveals a distinct, exterior KLA binding site. A) Cryo-EM reconstruction (3.6 Å) of MsbA(ADP·VO_4_) in complex with KLA. The density for KLA is shown in yellow, and MsbA subunits are shown in pink and blue. B) Coulombic electrostatic potential (scale bar -10 to +10 as computed by ChimeraX^50^) is colored red and blue for negative and positive charges, respectively. KLA and interacting residues are shown in ball and stick representation. C) MsbA shown with hydrophilic and hydrophobic surfaces colored blue and gold, respectively. D) Different views of KLA bound to MsbA. KLA and interacting residues shown in stick representation. Bonds are shown as dashed yellow lines. Residues are labelled. E) View of the bound ADP·VO_4_ and interacting residues shown as described in D.

### Probing the exterior KLA binding sites

A series of MsbA mutants engineered to impact KLA binding were evaluated. Some of the MsbA mutant proteins could be expressed and purified but were biochemically unstable, such as MsbA^Y87F,R238A^ (**Fig S19**). MsbA containing the R188A and K243A mutations (MsbA^R188A,K243A^), engineered to disrupt the interaction with one of the P-GlcN substituents of KLA, was biochemically stable and suitable for determining K_D_s (**Fig 5A-C and S20**). For the non-trapped protein, K_D1_ increased two-fold and K_D2_ increased five-fold. In the trapped state, K_D1_ and K_D2_ also both statistically increased by more than two-fold. As MsbA is known to be stimulated by some lipids^,22-24,^46 we determined the ATPase activity of MsbA in the absence and presence of lipid (**Fig 5D**). Wild-type MsbA was stimulated by KLA at levels observed by others^.17,^22 However, RaLPS (LRa), an LOS with a complete *E. coli* R2-type core,^47^ stimulated MsbA ATPase activity to a higher degree (**Fig 5D**).^22^ Consistent with a previous report,^22^ lipid A (LA), similar to KLA but lacking the Kdo substituents, stimulated MsbA but to lesser extent, highlighting the importance of the Kdo groups. MsbA^R188A,K243A^ displayed a statistically significant decrease in stimulation that was independent of the lipid type (**Fig 5D**). Lipid-induced stimulation of MsbA containing R78A and K299A mutations (MsbA^R78A,K299A^), engineered to disrupt binding at the interior site (**Fig S17B**), was also assessed. MsbA^R78A,K299A^ showed no stimulation by LA and KLA but activity was stimulated by LRa to the same level as the wild-type protein (**Fig 5D**). Next, we inspected the residues coordinating KLA in MsbA structures (**Fig 5E and S21**). Except for R188, the KLA interacting residues are in similar positions. However, in (occluded and open) OF states, TM4 is displaced ∼12 Å in a direction toward the other residues, priming R188 to engage the P-GlcN of KLA. This additional interaction explains the enhancement in KLA binding affinity. Together, these results indicate that the exterior KLA binding site has a direct role in allosterically stimulating MsbA ATPase activity.

**Figure 5.**
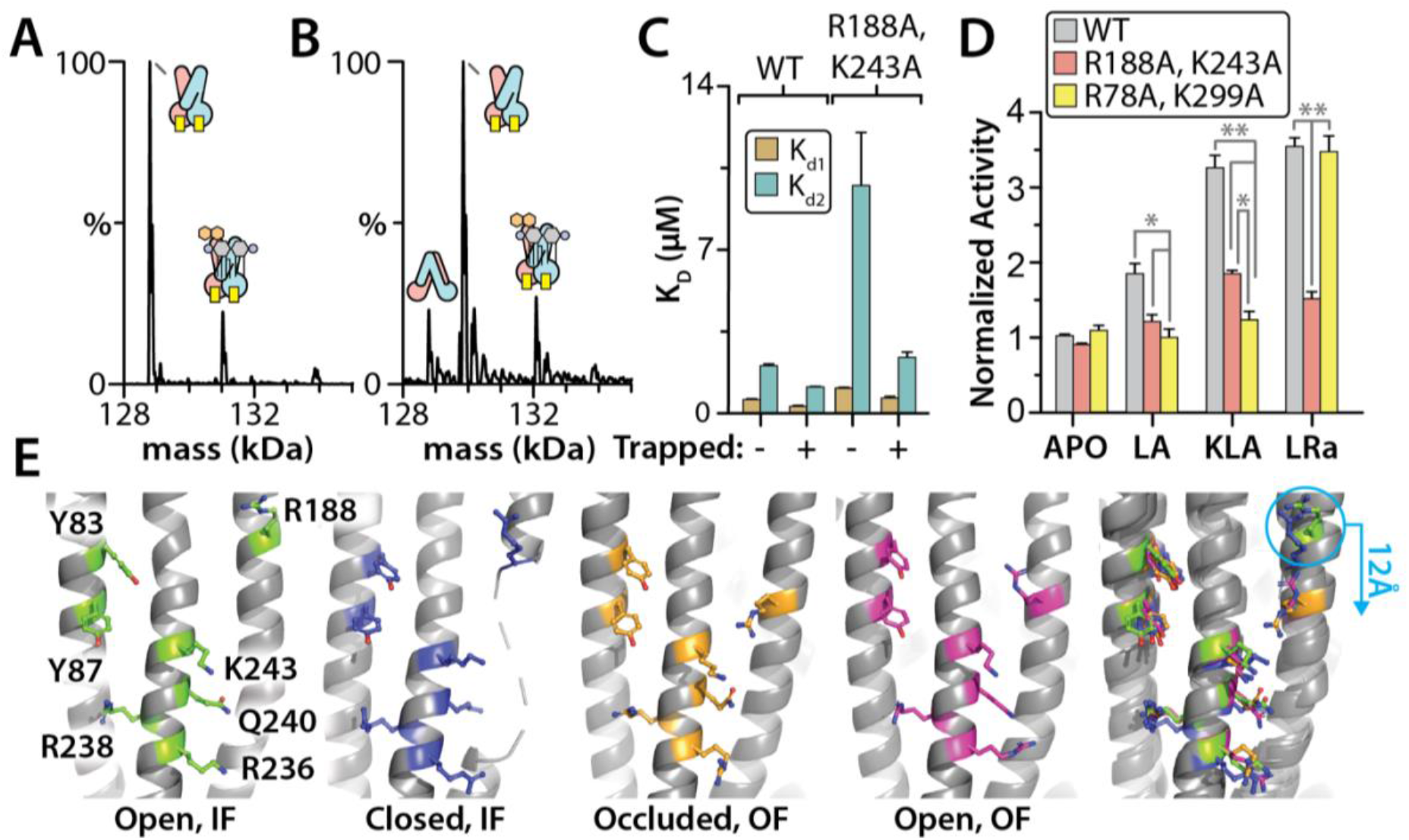
Characterization of the exterior KLA binding site. A) Deconvoluted mass spectrum of 0.3 *μ*M MsbA^R188A,K243A^ in the presence of 0.4 *μ*M KLA. B) Deconvoluted mass spectrum of 0.4 *μ*M vanadate-trapped MsbA^R188A,K243A^ with same concentration of KLA as in panel a. C) K_D_ values for KLA binding to the wild-type and mutant MsbA. D) ATPase activity of MsbA, MsbA^R188A,K243A^, and MsbA^R78A,K299A^ in the presence and absence of 5 *μ*M lipid A (LA), KLA, or Ra-LPS (LRa). E) Alignment of different structures to a region of TM5, residue range from 230 to 250. The first four panels shown (from left to right) correspond to apo (this report), G907 bound (PDB 6BPL), and vanadate-trapped in an occluded (PDB 7BCW) or open (this report) MsbA structures. The rightmost panel is an overlay of all four structures. Reported are the mean and standard deviation (*n* = 3). Student’s *t*-test (\**p* < 0.05, \*\**p* < 0.01).

### Implications of the exterior KLA binding site in MsbA

The unique KLA binding site uncovered here reveals an evolutionarily conserved feature of MsbA. The residues (R188, R238, and K243) that engage the characteristic P-GlcN substituents of LOS are conserved (**Fig S17D**). Similar interactions have been observed for the interior LOS site in MsbA^6,17^ and recognition of LPS in LptB2FG, an LPS ABC transporter.^48^ Other residues (R236 and Q240) that coordinate a Kdo group of KLA are also conserved (**Fig S17D**). As most LOS of gram-negative bacteria synthesize a KLA molecule resembling those found in *E. coli*,^3^ the conservation of residues and their interaction with KLA establish this as a unique feature of MsbA compared to other transporters. Moreover, the large, basic patch that nestles the KLA headgroup extends beyond the Kdo groups that could further engage the core oligosaccharide of LOS, the glycolipid MsbA flips. This basic patch likely extends to other MsbAs and important for recognition of LOS, including other lipids, that vary in structure in different bacteria.^2^

In summary, native mass spectrometry data reveal that MsbA co-purifies with copper(II), an observation that would remain unnoticed using traditional methods, and protein-lipid interactions are directly influenced by copper(II) binding as well as the conformation of the transporter. Structural studies show the N-terminus of MsbA coordinates copper(II) in a similar fashion as the GHK peptide. Copper(II) binding to MsbA impacts lipid binding, and the N-terminus is located near the bilayer where it plausibly interacts with lipid headgroups. More broadly, copper(II) binding may have a regulatory role, such as coupling copper(II) levels with LPS biogenesis, and warrants further investigation. Another structure illuminates a distinct, exterior KLA binding site is an allosteric modulator of ATPase activity and a conserved feature of MsbA. Mutagenesis studies document that this exterior, allosteric site is also sensitive to hexaacylated lipid A species, such as LA and LRa. These results provide compelling evidence of a feedforward activation mechanism for MsbA, a rare principal of control in biosynthetic pathways that is best described for pyruvate kinase,^49^ tuning MsbA activity to match cellular production of cytoplasmic LOS and precursors thereof.

## Acknowledgements

We thank Bryan Tomlin in the Elemental Analysis Laboratory for elemental analysis, Jonathan Schuermann and Igor Kourinov at APS for useful discussion regarding data collection. Part of this work is based upon research conducted at the Northeastern Collaborative Access Team beamlines (P30 GM124165) at the Advanced Photon Source (DE-AC02-06CH11357). This work was also supported by National Institutes of Health (NIH) under grant numbers (DP2GM123486, R01GM121751, R01GM139876 and R01GM138863 to A.L.; R35GM143052 to M.Z.; and P41GM128577 to D.R.). We thank the staff at the University of Chicago Advanced Electron Microscopy (RRID:SCR_019198) for the help with cryo-EM data collection. We thank Research Computing Center at the University of Chicago for the support of this work by providing the computing resources of the Beagle3 HPC cluster funded by NIH (S10OD028655).

## Author Contributions

J.L. and A.L. designed the research. J.L. and T.Z. expressed and purified MsbA. J.L. performed mass spectrometry experiments. J.L. and T.Z. carried out the functional assays. J.L. and A.L. analyzed the data. C.L. and M.Z. collected and processed cryo-EM data. S.S., J.L., T.Z., and A.L. performed X-ray crystallography. S.S., C.L., S.S., J.L., M.Z., and A.L. built the atomic models. G.H. and A.L. analyzed MsbA sequences. J.L. and A.L. wrote the manuscript with input from the other authors.

## Data availability

Atomic coordinates and structure factors for the crystal structure of an N-terminal fragment of MsbA fused to GFP and bound to copper(II) has been deposited in the Protein Data Bank (PDB) under accession code 8DHY. The coordinates for apo and KLA-bound MsbA cryoEM structures have been deposited in the PDB under accession codes and 8DMO (EMD-27545) and 8DMM (EMD-27544), respectively.

## Materials & Correspondence

Correspondence and requests for materials should be addressed to AL.

