## Supplementary Information for "Structural basis for lipid and copper regulation of the ABC transporter MsbA"

### Materials and Methods

***Construction of MsbA and other expression plasmids.*** The MsbA gene and pCDF-1b plasmid (Novagen) were amplified by polymerase chain reaction (PCR) using Q5 High-Fidelity DNA Polymerase (New England Biolabs, NEB) from *Escherichia coli* genomic DNA and purified plasmid, respectively. Primers were designed using the online NEBuilder Assembly Tool (NEB) and amplified products were gel purified prior to HiFi DNA Assembly (NEB) following the manufacturer's protocol. The resulting construct, pCDF-MsbA, expressed MsbA with an N-terminal TEV cleavable $His_6$ fusion protein. To generate mutant forms of MsbA, primers were designed using online tool NEBaseChanger (NEB) and mutants introduced using the KLD enzyme mix (NEB) following the manufacturers protocol. The N-terminal sequence of MsbA was grafted onto MBP, superfolder GFP[1] and T4 lysozyme by subcloning into the pCDF-MsbA plasmid and keeping residues 1-8 of MsbA. A similar fusion strategy has been done for the GHK peptide.[2] Truncation of the MsbA N-terminus was carried using KLD enzyme mix (NEB) following manufacturers protocol. The GFP fusion for successful structure determination had a N-terminal sequence after TEV protease cleavage of MHNDKGEELF with the MsbA sequence underlined. All plasmids were confirmed by DNA sequencing.

***MsbA expression and membrane preparation.*** The wild-type and mutant MsbA expression plasmids were transformed into *E. coli* (DE3) BL21-AI competent cells (Invitrogen) and incubated at 37 °C until the $OD_{600nm}$ ≈ 0.6-1.0 at which point the cultures were induced with final concentrations of 0.5 mM IPTG (isopropyl β-D-1-thiogalactopryanoside) and 0.2% (w/v) arabinose. The cultures were induced overnight at 25 °C. The cultures were then harvested at

4500 x g for 12 minutes and the resulting pellet was resuspended in 20 mM Tris, 300 mM NaCl, pH 7.4 and supplemented with Roche cOmplete Protease Inhibitor Cocktail tablet. The suspension was lysed in a Microfluidics M-110P microfluidizer operating at 25,000 psi on ice. The lysate was centrifuged at 40,000 x g for 20 minutes and the resulting supernatant was centrifuged at 100,000 x g for 2 hours. The resulting pellets were collected and homogenized in 20 mM Tris, 150 mM NaCl, 20% (v/v) glycerol, pH 7.4. The membrane solution was extracted with 1% (w/v) DDM, rotating overnight at 4 °C. The extraction was then centrifuged at 40,000 x g for 10 minutes and the resulting supernatant was supplemented with 10 mM imidazole and filtered with a 0.45 µm syringe filter.

***Detergent screening and optimization of purification.*** The extracted material was subjected to extensive detergent screening[3] to determine the delipidating ability of each detergent on MsbA. In short, His-tagged MsbA was bound to 100 µL Ni-NTA beads (Qiagen) equilibrated with NHA buffer (20 mM Tris, 150 mM NaCl, 10 mM imidazole, 10% (v/v) glycerol, pH 7.4) supplemented with 2x the critical micelle concentration (CMC) of DDM and then washed with 5 column volumes (CV) of NHA containing 2x CMC DDM buffer. The bound protein was then treated with 10 CV of NHA containing 2x CMC DDM buffers supplemented with 10x CMC of various detergents (Anatrace). The column was then re-equilibrated with 5 CV of NHA-2x CMC DDM buffer and eluted with 2 CV of NHA containing 2x CMC DDM buffer supplemented with 500 mM imidazole. To check the degree of delipidation via native mass spectrometry, the eluent was buffer exchanged into 200 mM ammonium acetate, 2x CMC DDM, pH 7.4 via Micro Bio-Spin P-6 gel centrifuge columns (Biorad) following the manufacturer's protocol. After determination and purification with the optimal detergent wash (NG), the protein was buffer exchanged back into NHA-2x CMC DDM on a HiPrep 26/10 desalting column (GE Healthcare). The sample was then treated with TEV protease, produced in-house, overnight at room temperature to remove the N-terminal His tag, 10mM β-mercaptoethanol was added during the TEV treatment. The digested material was passed over Ni-NTA agarose equilibrated with NHA-2x CMC DDM and the flow-through containing the cleaved material was collected. The material was concentrated using a centrifugal concentrator (Millipore, 100 kDa molecular weight cutoff) followed by injection onto a Superdex 200 Increase 10/300 GL (GE Healthcare) column equilibrated with 20 mM HEPES, 200 mM NaCl, 10% (v/v) glycerol and 2x CMC $C_{10}E_5$. Peak fractions containing dimeric MsbA were pooled and flash frozen at -80 °C.

**Preparation of MsbA for native MS and structural studies.** For MsbA with reduced copper(II) binding, additional 1mM $MgCl_2$ was added to all buffer and MsbA solution in the purification step using Ni-NTA beads. Copper(II) saturated MsbA was obtained by adding 20uM copper (II) acetate then buffer exchanged using Bio-Spin column to remove excess copper(II). MsbA trapped by vanadate was obtained by adding ATP and $MgCl_2$ to MsbA to reach the final concentration of 10mM for both then incubating at room temperature for 10min. After incubation, vanadate (pH 10) was added to reach final concentration of 1uM followed by incubation at 37°C for 10min. MsbA samples was buffer exchanged using Bio-Spin column to 200mM ammonium acetate supplemented with 2x CMC $C_{10}E_5$ for native MS studies. To prepare MsbA samples for Cryo-EM studies, MsbA and trapped MsbA were preloaded with copper(II) then purified by Superdex 200 Increase 10/300 GL size exclusion column equilibrated with 200mM NaCl, 20mM HEPES and 2x

CMC $C_{10}E_5$ without glycerol. Peak fractions containing MsbA were pooled and concentrated to 8mg/mL then mixed with KLA at 1:2 molar ratio (1 KLA for 1 MsbA subunit).

***Native Mass Spectrometry.*** Samples were loaded into gold-coated glass capillaries made in-house[3] and were ionized via electrospray into a Thermo Scientific Exactive Plus Orbitrap with Extended Mass Range (EMR). For native mass analysis, the instrument was tuned as follow: source DC offset of 25, injection flatapole DC to 8.0 V, inter flatapole lens to 7, bent flatapole DC to 6.0, transfer multipole DC to 2 and C trap entrance lens to 2, trapping gas pressure to 6.0 with the in-source CID to 60.0 eV and CE to 100, spray voltage to 1.70 kV, capillary temperature to 200 ˚C, maximum inject time to 200 ms. Mass spectra were acquired with setting of 17,500 resolution, microscans of 1 and averaging of 100.

***Determination MsbA-lipid equilibrium binding constants.*** Lipids were prepared as previously described,[4] in which lipids dissolved in chloroform were dried under nitrogen flow placed under vacuum overnight followed by dissolving in water. The concentration of MsbA was determined using a DC protein assay (BioRad) with bovine serum albumin as the standard. MsbA was incubated with lipids at varying concentrations and mixed with 200mM ammonium acetate supplemented with 2x CMC lauryldimethylamine oxide (LDAO) at 1:1 volumn ratio. Samples were incubated in the nano electrospray ionization source chamber for a minute to reach equilibrium prior to data acquisition. These samples analyzed on Orbitrap Exactive Plus EMR mass spectrometer (Thermo Scientific) operating at identical settings as described above. The mass spectra for each titration event were obtained in triplicate. The mass spectra were deconvoluted using UniDec[5] and the resulting peak intensities for apo and lipid-bound protein were determined. The relative abundance for each species was determined by dividing the peak intensity by the total intensity to convert to mole fraction for each independent experiment. For MsbA (P) binding the $N^{th}$ lipid ($L_n$), we applied the following sequential lipid binding model:

$$PL_{n-1} + L \overset{K_A}{\Leftrightarrow} PL_n$$

where:

$$K_{An} = \frac{[PL_n]}{[PL_{n-1}][L]}$$

To calculate the mole fraction of a particular species:[4]

$$F_{PLn} = \frac{[L]_{free}^{n} \prod_{j=1}^{n} K_{Aj}}{1 + \sum_{i=1}^{n} [L]_{free}^{i} \prod_{j=1}^{n} K_{Aj}}$$

For each titrant in the titration the free concentration of lipid was computed as follows:

$$[L]_{free} = [L]_{total} - [P]_{total} \sum_{i=0}^{n} iF_{PLi}$$

The sequential lipid binding model was globally fit to the mole fraction data by minimization of pseudo-$\chi^2$ function:

$$\chi^2 = \sum_{j=1}^{m} \sum_{k=1}^{d} (F_{i,j,exp} - F_{i,j,calc})^2$$

where *n* is the number of bound ligands and *d* is the number of the experimental mole fraction data points.

***Determination of metal bound to MsbA.*** Samples of MsbA were submitted to the Elemental Analysis Laboratory at Texas A&M University for elemental analysis by inductively coupled plasma mass spectrometry. A NexION ICP mass spectrometer (PerkinElmer) was used with operating parameters provided in Table S2.

***MsbA Activity Assay.*** The ATPase activity of MsbA was determined following a modified version of the malachite green assay.[6] For the fully delipidated (apo) protein, 400 nM of MsbA (in 200 mM ammonium acetate supplemented with 1x CMC $C_{10}E_5$ and 1x CMC LDAO) was incubated with 5 mM $MgCl_2$ and 200 µM ATP at 37 °C for 12 minutes. For analysis of protein with KLA, lipid was added to the sample at a final concentration of 5 µM. Endpoint samples were collected at 3, 6, 9, 12 minutes and stopped with the addition of malachite green solution with the following components: 3:1 mixture of 0.045% (w/v) malachite green and 4.2% (w/v) ammonium molybdate prepared in 4 N HCl, 0.04% (v/v) Triton X-100 (final concentration). Then 34% (w/v) sodium citrate was added to stop the coloring reaction. The quenched reactions were incubated at room temperature for 30 minutes and absorbance at 650 nm was measured on a CLARIOstar plate reader (BMG LabTech). The hydrolyzing rate of ATP was obtained by plotting the slope of absorbance of samples collected at 3, 6, 9, and 12 minutes.

***X-ray structure of N-terminal peptide bound to copper(II).*** Initial crystallization trials were carried out for the GFP fusion protein at a concentration of 1 mM (using an extinction coefficient at 490 nm of 39.2 x $10^3$ $M^{-1}cm^{-1}$)[1] using a Mosquito LCP (TTP Labtech) crystallization robot in hanging drop plates at 20°C. Crystals grew in index condition C5 (60% Tacsimate pH 7.0) and were further optimized by increasing the concentration of Tacsimate pH 7.0 to 70%. Crystals were cryoprotected using 100% Tacsimate pH 7.0. Single crystals were mounted with CrystalCap HT

Cryoloops (Hampton Research) prior to flash freezing in liquid nitrogen. The initial diffraction data were collected in-house on a Rigaku Raxis-IV++. Initial phases were determined using molecular replacement with PDB code 2B3P. Model refinement and building were performed using Phenix[7] and Coot.[8] Anomalous data was collected using a wavelength of 1.378Å at the Advanced Photon Source on beamline 24-ID-C. Although the structure could be determined by SAD phasing using Phenix AutoSol program, the model built from the in-house data was used for molecular replacement followed by model building and refinement.

***Sample preparation for single-particle cryo-EM.*** Vitrification was performed using a Vitrobot Mark IV (Thermo Fisher) operating at 8 °C and 100% humidity. A total of 3.5 µL of sample (8 mg/mL copper-loaded MsbA either apo or trapped with ADP-vanadate in 200mM NaCl, 20mM HEPES, supplemented with 2x CMC $C_{10}E_5$) was applied to holey carbon grids (Quantifoil 300 mesh Cu 1.2/1.3) glow-discharged for 30 seconds. Samples contained two-fold molar excess of KLA. The grids were blotted for 5 seconds at blotting force 1 using standard Vitrobot filter paper (Ted Pella, 47000-100), and then plunged into liquid ethane.

***Data collection for single-particle cryo-EM.*** The optimized grids were sent to the Advanced Electron Microscopy Facility at the University of Chicago for data collection. The dataset was collected as movie stacks with a Titan Krios electron microscope operating at 300 kV, equipped with a K3 direct detector camera. Images were recorded at a nominal magnification of 81,000x at super-resolution counting mode by image shift. The total exposure time was set to 4 s with a frame recorded every 0.1 s, resulting in 40 frames in a single stack with a total exposure around

50 electrons/Å$^2$. The defocus range was set at -1.0 to -2.5 µm. See Table S8 for the details of data collection parameters.

***Image processing for single-particle cryo-EM.*** Collected movies were subjected to motion correction by MotionCor2.[9] Subsequent processing was carried out in cryoSPARC.[10] The detailed data processing flow is shown in Fig S15 (vanadate-trapped MsbA) and Fig S22 (open, inward-facing MsbA). Stage drift and anisotropic motion of the stack images were first corrected by patch-based motion correction. CTF parameters for each micrograph were determined by patch-based CTF estimation. For vanadate-trapped MsbA, the particles were picked using the templates generated from the blob picker. For the open, inward-facing MsbA, the final particle set was picked using templates generated from a 3D model from an earlier reconstruction. For both datasets, the particles were cleaned by two rounds of 2D classification. Three initial models were generated from the remaining particles using ab initio reconstruction. The particles were further classified by heterogeneous refinement based on three initial models. For the vanadate-trapped MsbA, the best class of particles was selected for a non-uniform refinement with per-particle defocus and CTF optimization, and a C2 symmetry imposed, resulting in a final map resolved at 3.6 Å. For the open MsbA, the best class of particles was selected for a non-uniform refinement with either C2 or C1 symmetry imposed, resulting in final maps resolved at 3.88 Å and 4.06 Å, respectively. See Table S8 for the details of image processing statistics.

***Model building, refinement, and validation for single-particle cryo-EM structures.*** The previously reported structure[11] of MsbA with ADP-vanadate from *Escherichia coli* (PDB 5TTP) was docked into the cryo-EM map using Chimera.[12] The model was manually refined using Coot.[8] Phenix[7] was used to generate the coordinates and restraint files for KLA. The final model underwent one round of real-space refinement using Phenix with secondary-structure and Ramachandran restraints. Geometry outliers were manually fixed in Coot. The statistics of the final round of model refinement and the model geometry are reported in Table S9. Figures were generated using ChimeraX[13] and Pymol (Schrödinger LLC., version 2.1). See Table S9 for the details of model statistics.

***Conservation and sequence analysis of bacterial MsbA.*** Sequences for conservation analysis were gathered using NCBI Blast and the *E. coli* MSBA protein as input. We separately searched for MSBA sequences from gammaproteobacteria, deltaproteobacteria, alphaproteobacteria, epsilonproteobacteria, the FCB clade, and Bacilli to get broad representation across Bacteria. We then aligned the sequences with Muscle 3.83. We removed all gaps caused by sequence absent in the *E. coli* sequence and extracted sites interfacing with KLA using a custom python script. A subalignment containing only these sites was used with the weblogo server (https://weblogo.berkeley.edu/) to generate the sequence logo. For analysis of N-terminal sequences of ABC transporters, more than >20k MsbA sequences were downloaded from UniProt. A python script making use of BioPython[14] analyzed sequences containing an N-terminal sequence of to begin with MH.

**Supplementary Text**

***Optimization of MsbA samples.*** An established detergent screening method was employed to optimize the purification of MsbA.[3] In brief, MsbA bound to nickel affinity resin was subsequently washed with various detergents. Of the detergents screened, those with short acyl chains (OG: n-octyl-β-D-glucopyranoside; OGNG: octyl glucose neopentyl glycol; LMNG: lauryl maltose neopentyl glycol) had the greatest impact on MsbA homogeneity. For example, the mass spectrum for MsbA washed with OG revealed mass spectral peaks corresponding to dimeric MsbA. In addition, the mass spectrum also indicates the presence of several adducts including a second charge state distribution that is ~3.5 kDa heavier in molecular weight. The adduct mass agrees with the mass of LPS, a lipid we found co-purifying with the mechanosensitive channel of large conductance.[15] LPS has been reported to co-purify with the transporter.[11,16] Pure samples of MsbA, devoid of contaminating lipids, was readily achieved by washing with n-nonyl-β-D-glucopyranoside (NG) followed by purification via size exclusion chromatography in the charge-reducing detergent pentaethylene glycol monodecyl ether ($C_{10}E_5$).[3,17] The use of charge-reducing detergents aids preservation of non-covalent interactions and native-like structure in the mass spectrometer.[17,18] These results indicate commonly prepared MsbA samples are heterogeneous, which poses significant challenges for employing high-resolution native MS to interrogate membrane protein-lipid interactions.

***CryoEM structure of open, inward-facing MsbA.*** MsbA in the presence of two equivalents of KLA was subjected to cryo-electron microscopy (cryoEM) studies. The structure of MsbA was determined to a resolution of 3.9 Å (**Fig S22-S23 and Table S8**). The maps were of sufficient quality to build the TMD. However, density in some regions of the NBDs was less clear. MsbA adopts an open, inward facing conformation with the NDBs separated by ~57 Å from the C$\alpha$ of R569 of one subunit to the other. In addition, tube-like density is observed in the central cavity and on outside, membrane facing side near residue 87 (**Fig S23b**)**.** However, density was not clear enough to place the lipid.

**Supplementary Figures**

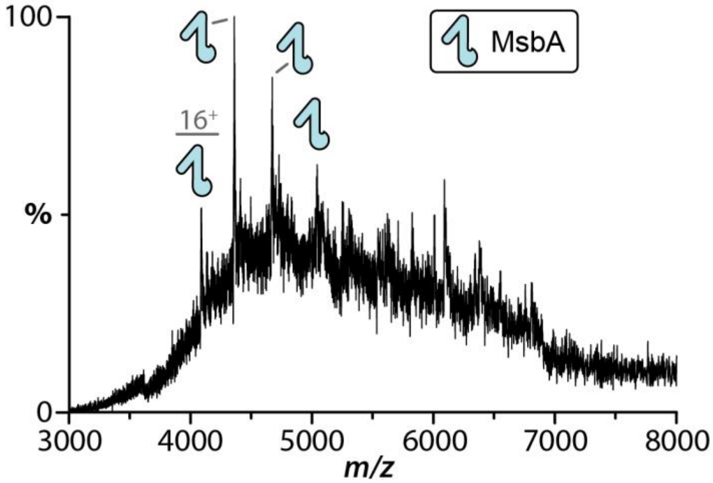


**Figure S1. Mass spectrum of MsbA purified in DDM following methods used for structural studies.** The underlying broad hump indicates the sample is highly heterogeneous. Peaks corresponding to monomeric MsbA are result of dissociation of complex under the high energy instrument settings.

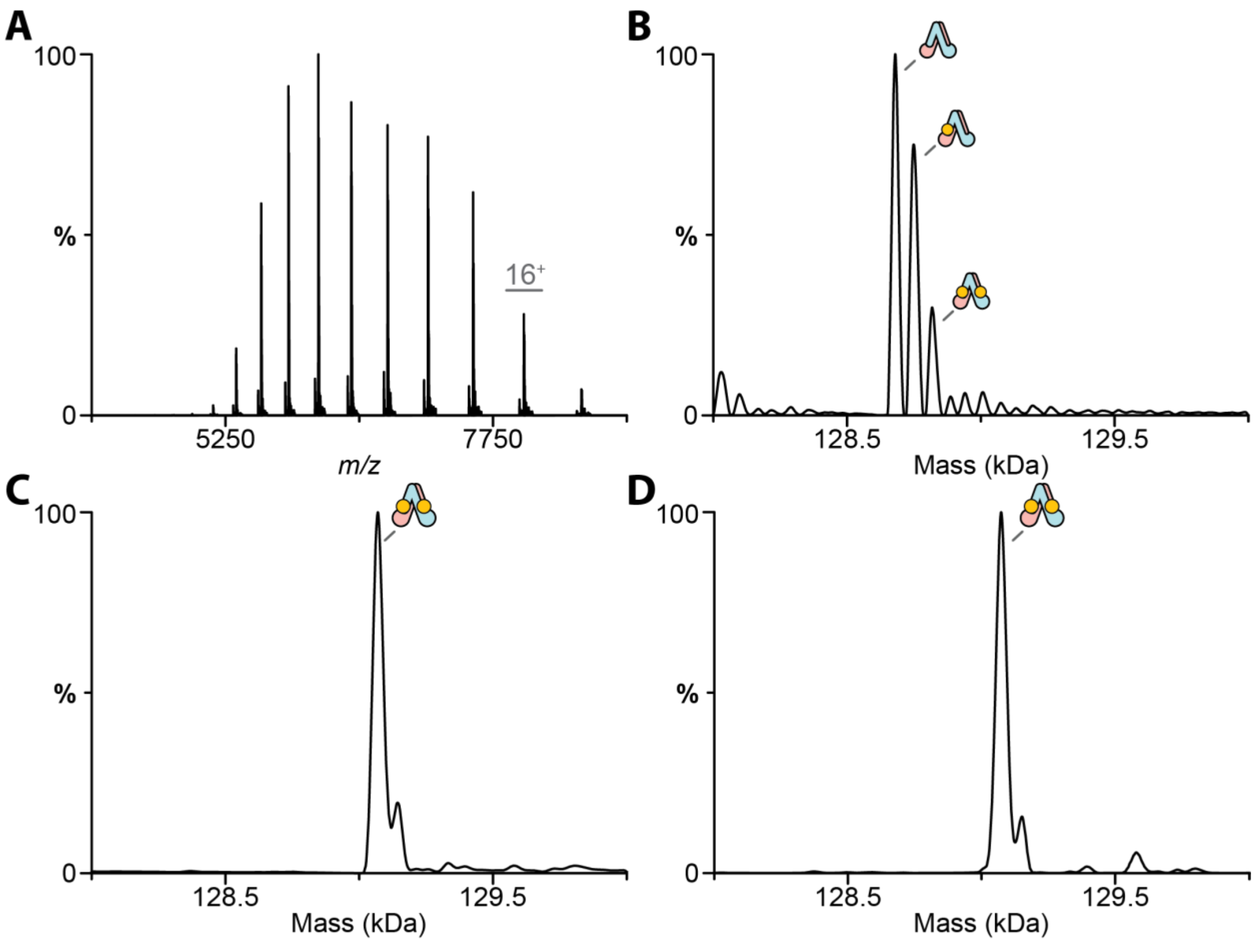


**Figure S2. Copper(II) binding to mutant MsbA(H562A, H576A) and ineffective chelation of copper(II) with trientine.** A) Native mass spectrum of $MsbA^{H562A,H576A}$ as isolated. B) Deconvolution of the mass spectrum shown in panel. Up to two copper(II) ions are bound to the mutant transporter. C-D) Deconvolution of wild-type MsbA after C) loading with copper(II) and D) after incubation with 2mM trientine. The specific chelator is unable to remove the MsbA bound copper(II).

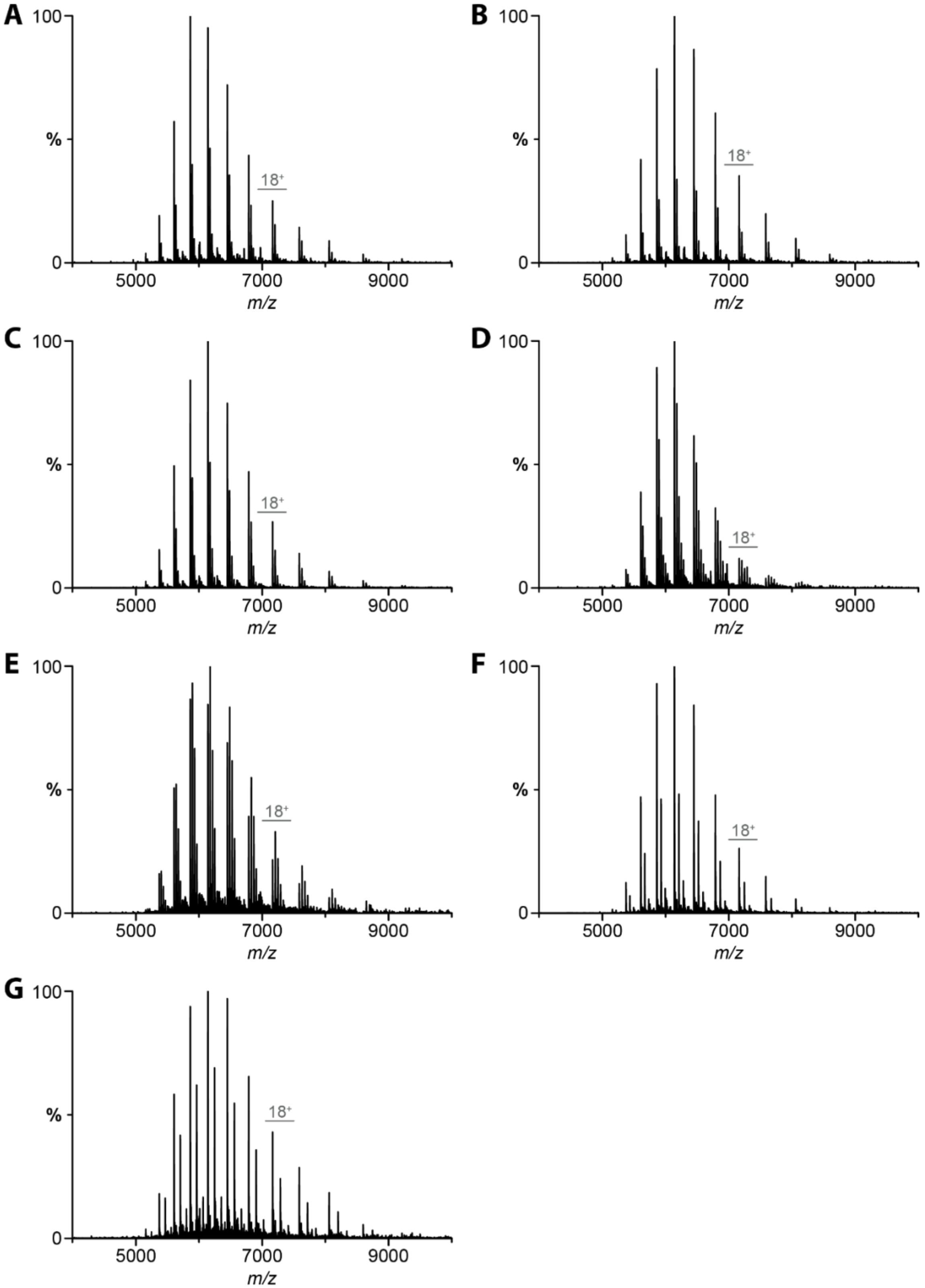


**Figure S3. Representative native mass spectra for lipid binding to MsbA partially loaded with copper(II).** MsbA (0.5 µM) was mixed with 6 µM of A) POPA, B) POPC, C) POPE, D) POPG, or E) POPS and 1µM of F) TOCDL, G) KLA.

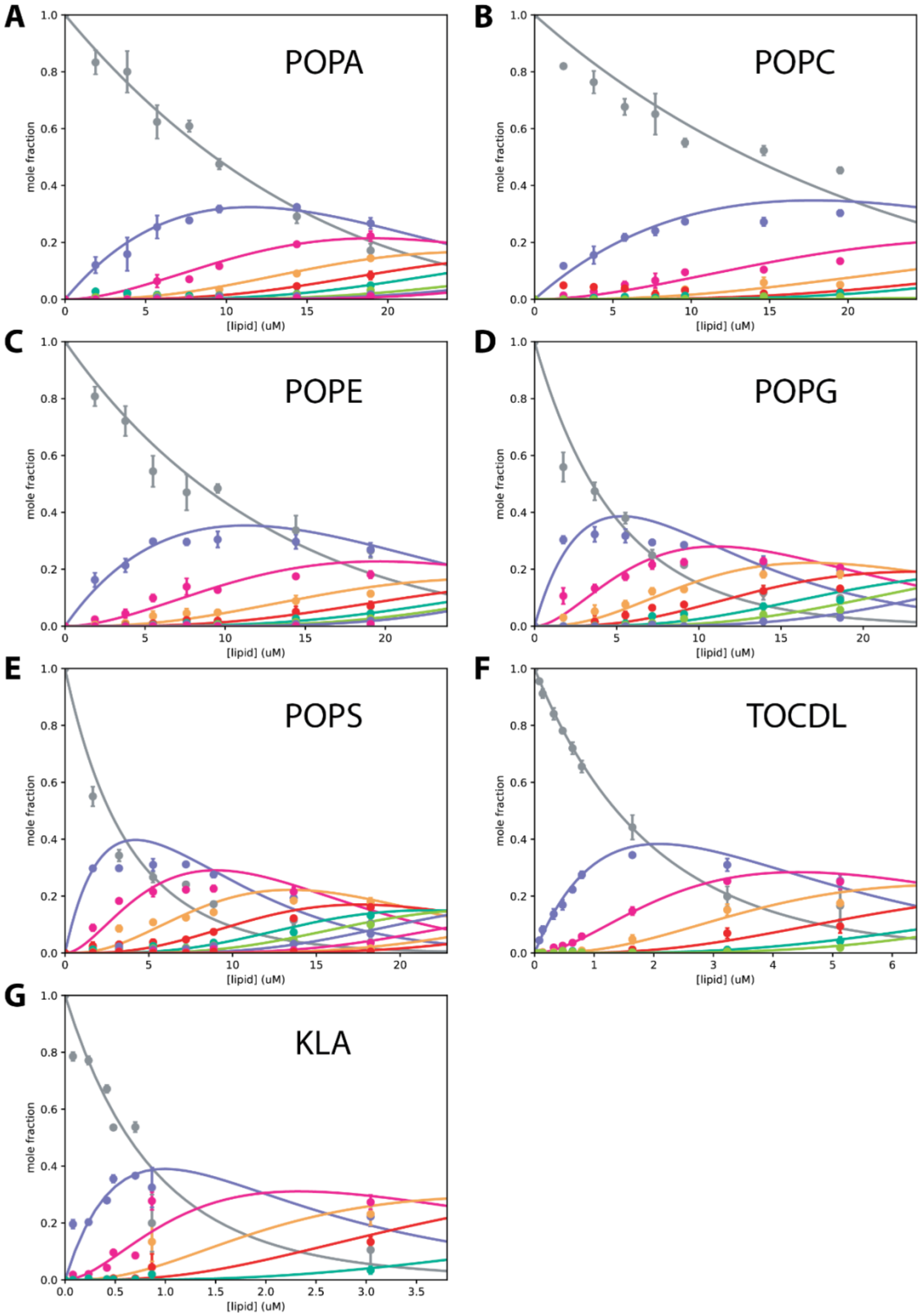


**Figure S4. Determination of equilibrium dissociation constants ($K_D$) for lipids binding MsbA partially loaded with copper(II).** Plots of mole fraction for apo MsbA bound to different number of lipids (dots) and resulting fit from a sequential lipid-binding model (solid lines). The different lipids are labelled.

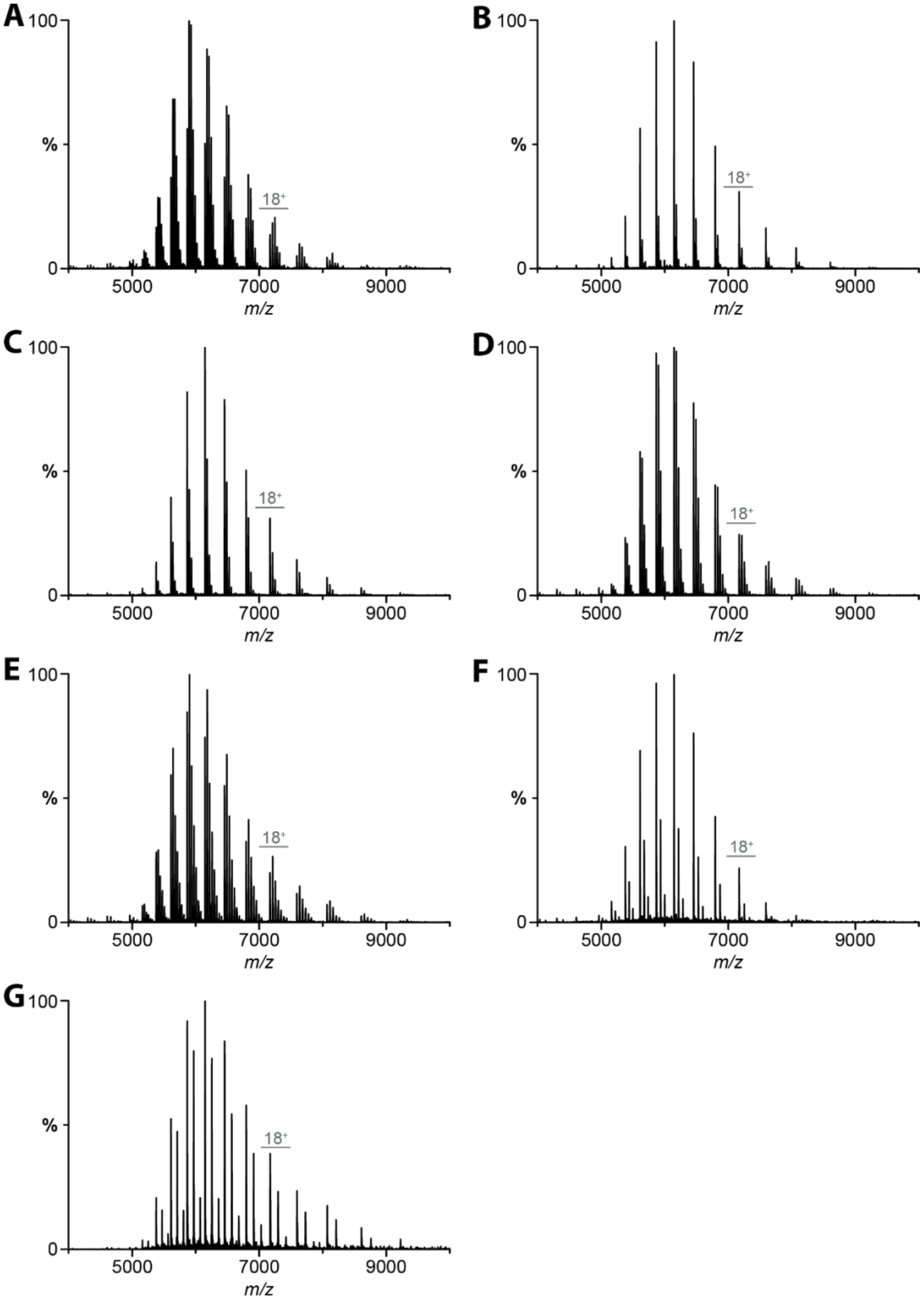


**Figure S5. Representative native mass spectra for lipid binding to MsbA fully loaded with copper(II).** MsbA (0.5 µM) was mixed with 6 µM of A) POPA, B) POPC, C) POPE, D) POPG, E) POPS and 1µM of F) TOCDL. G) 0.43 µM. For KLA, the concentration is 0.7 µM of KLA.

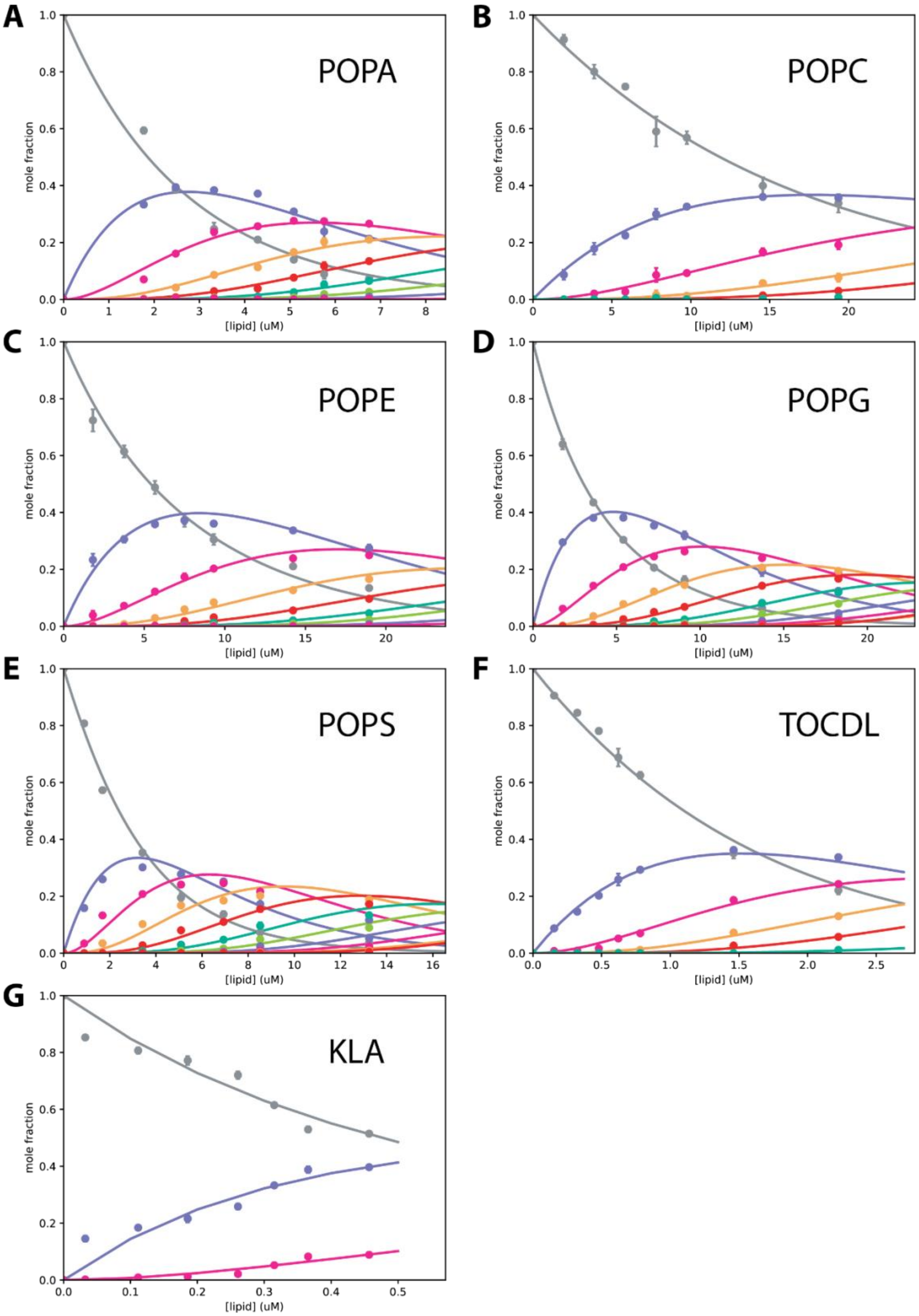


**Figure S6. Determination of equilibrium dissociation constants ($K_D$) for lipids binding MsbA fully loaded with copper(II).** Shown as described in Figure S4.

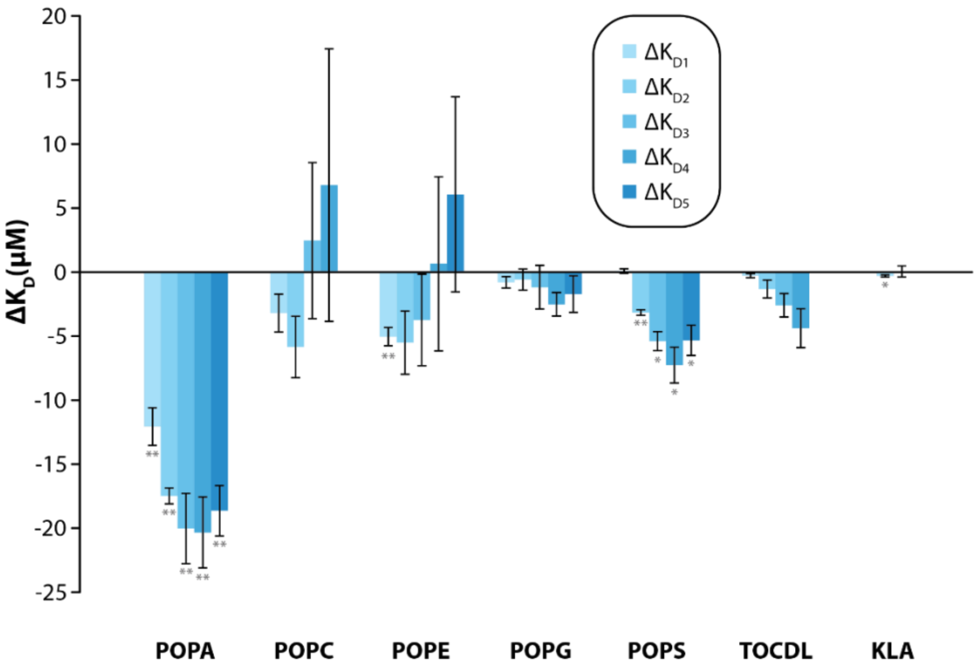


**Figure S7. Copper binding impacts MsbA-lipid interactions.** Reported are the difference $K_D$ values for MsbA saturated with copper(II) minus partially loaded. The asteriks denote student's t-test for MsbA partially versus fully loaded with copper(II) (**p*<0.05, ***p*<0.01)

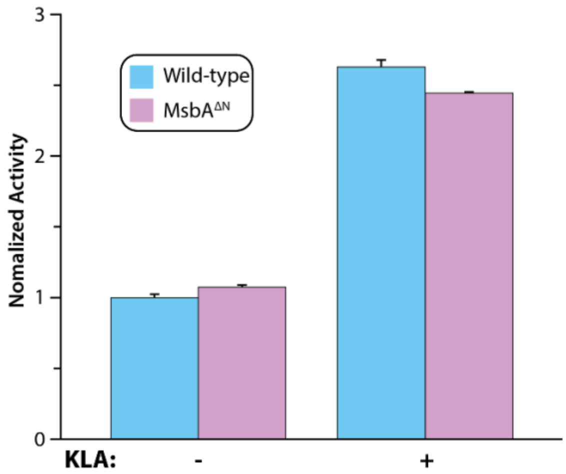


**Figure S8. ATPase activity for wild-type and N-terminal truncated MsbA.** KLA was added to a final concentration of 5 μM. Reported are the mean and standard deviation (*n* = 3).

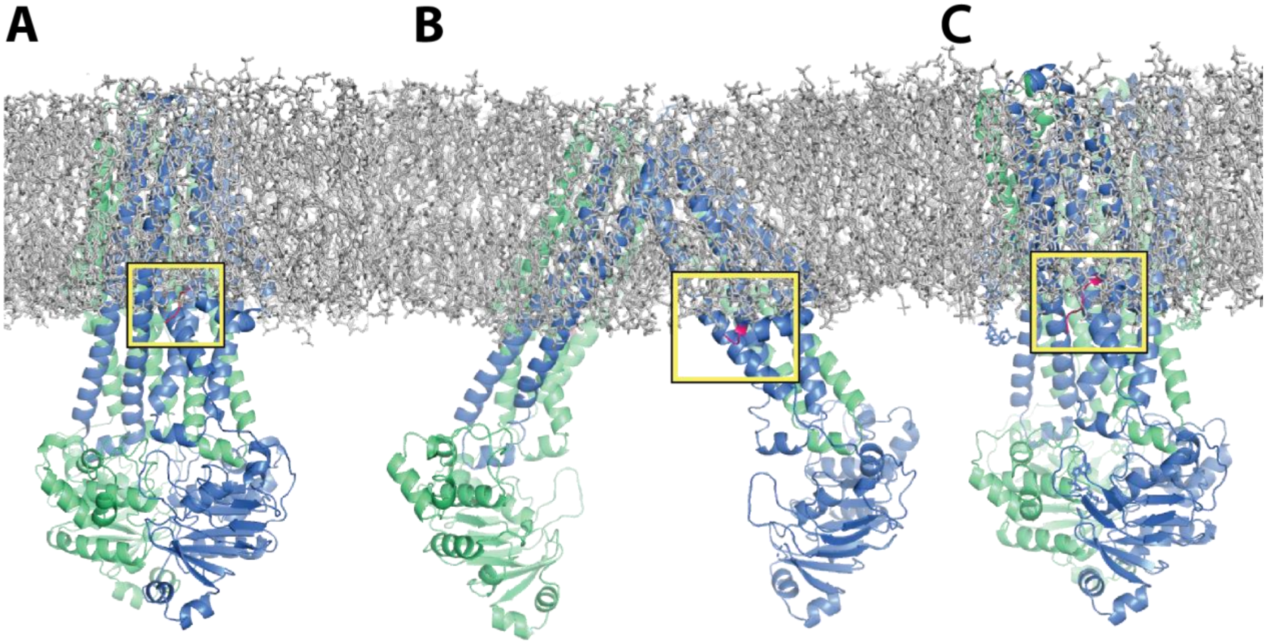


**Figure S9. N-terminal copper(II) binding site is located in proximity of the bilayer.** A-C) Snapshots from molecular dynamics simulation of MsbA structures in 16:0 PC (DPPC) bilayer (PDB 6BPP downloaded from MemProtMD.[19] DPPC lipids are shown in grey sticks, and MsbA in cartoon representation. Part of the N-terminus (first residue in model to residue 9) in colored pink. The yellow box highlights the position of the N-terminus relative to the bilayer. The coordinate files downloaded from MemProtMD are A) 7BCW structure is MSBA in salipro with ADP vanadate, B) open, inward-facing 6BL6 with protein replaced by open, inward-facing MsbA (this report), and C) 3B60 with protein replaced by MSBA trapped with ADP and vanadate.

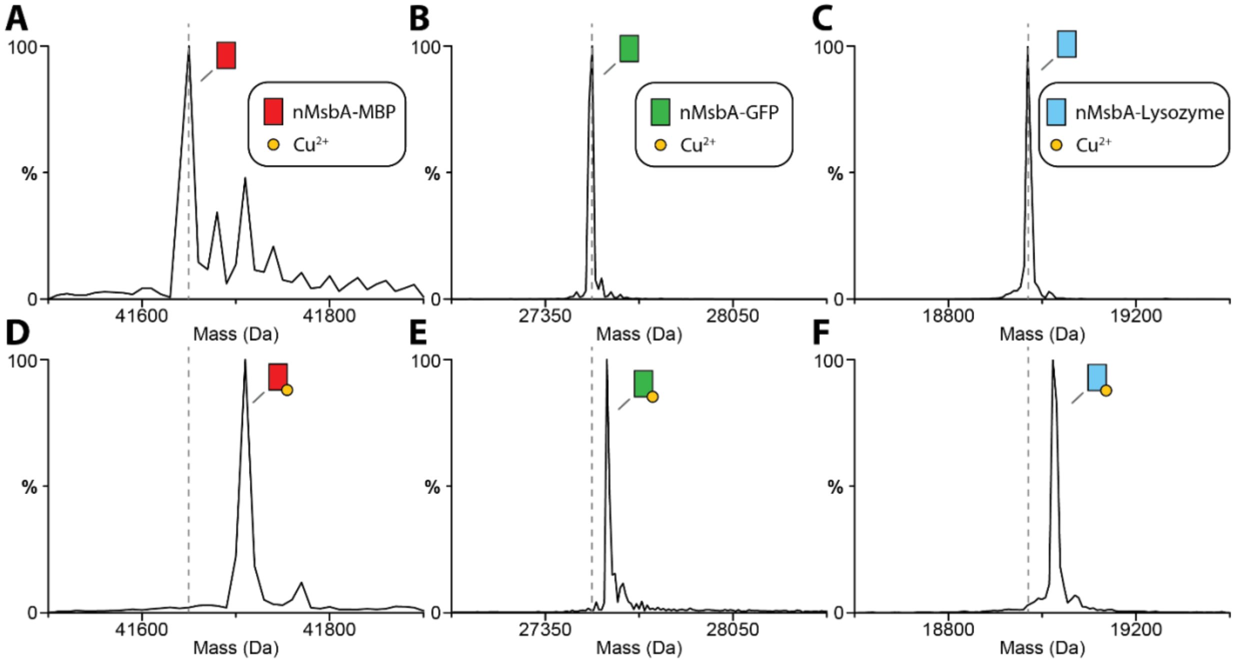


**Figure S10. Fusion proteins containing a fragment of the N-terminal sequence from MsbA bind copper(II).** Shown are the decovoluted mass spectra of A-C) MBP, GFP and T4 lysozyme before addition of copper(II). D-F) MBP, GFP and lysozyme after addition of copper(II).

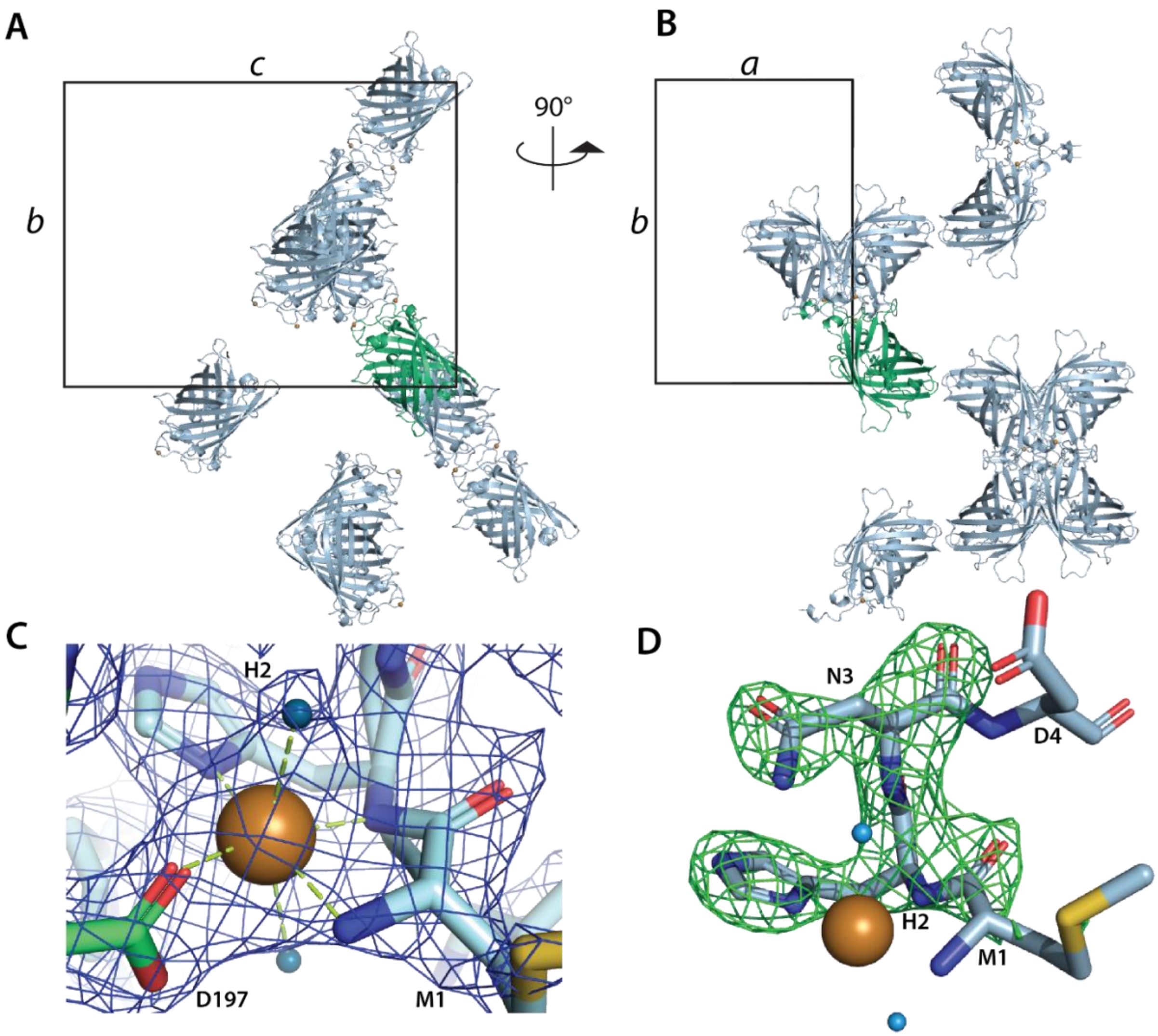


**Figure S11. Crystal structure of the N-terminal sequence of MsbA fused to GFP and bound to copper(II).** A-B) Crystal packing with one molecule (green) in the asymmetric unit cell and symmetry related molecules shown in light blue. C) Coordination of copper(II) within the crystal lattice. D197' from a symmetry related molecule coordinates copper(II). $2F_o–F_c$ electron density map contoured at 1.5 sigma is shown. d) Electron difference density map ($F_o$-$F_c$) after refinement with the N-terminus (residues 1-3) omitted and contoured at 5 sigma.

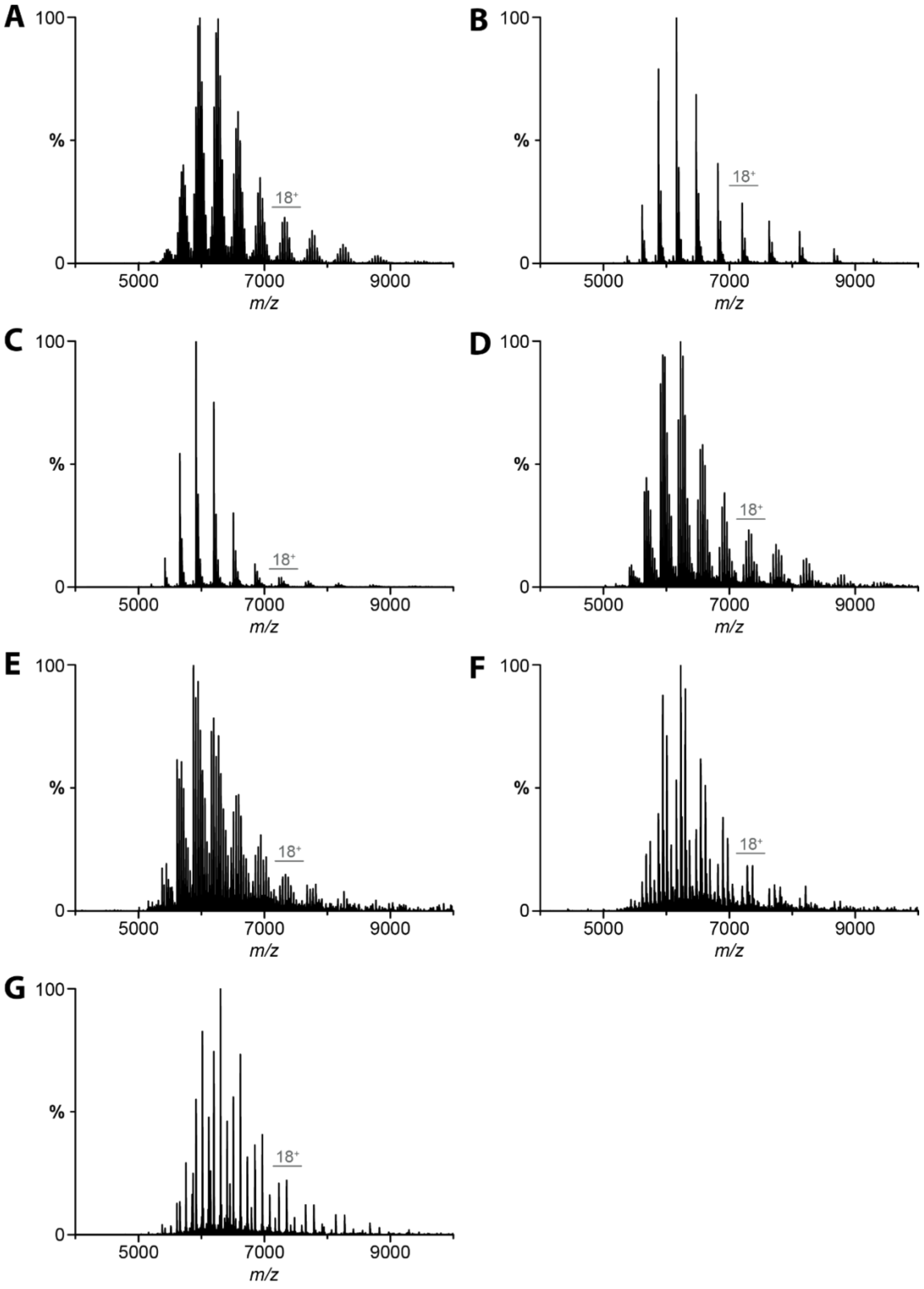


**Figure S12. Representative native mass spectra for lipids binding to MsbA trapped with ADP and vanadate.** MsbA (0.36 μM) was mixed with 6 μM of A) POPA, B) POPC, C) POPE, D) POPG, E) POPS and 1μM of F) TOCDL. G) 0.45 μM MsbA was mixed with 0.7 μM of KLA.

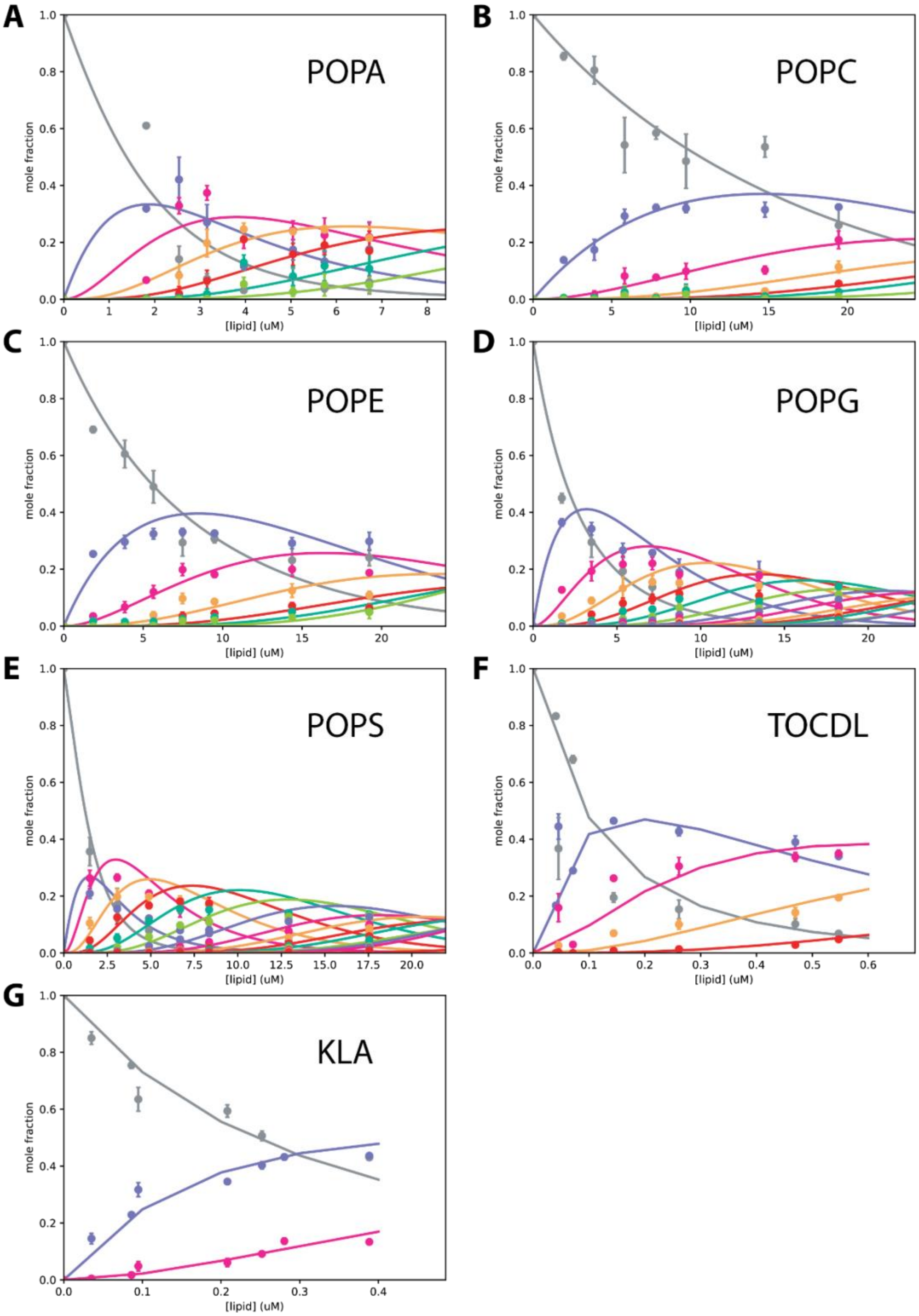


**Figure S13. Determination of equilibrium dissociation constants ($K_D$) for lipids binding MsbA trapped with ADP and vanadate.** Shown as described in Figure S4.

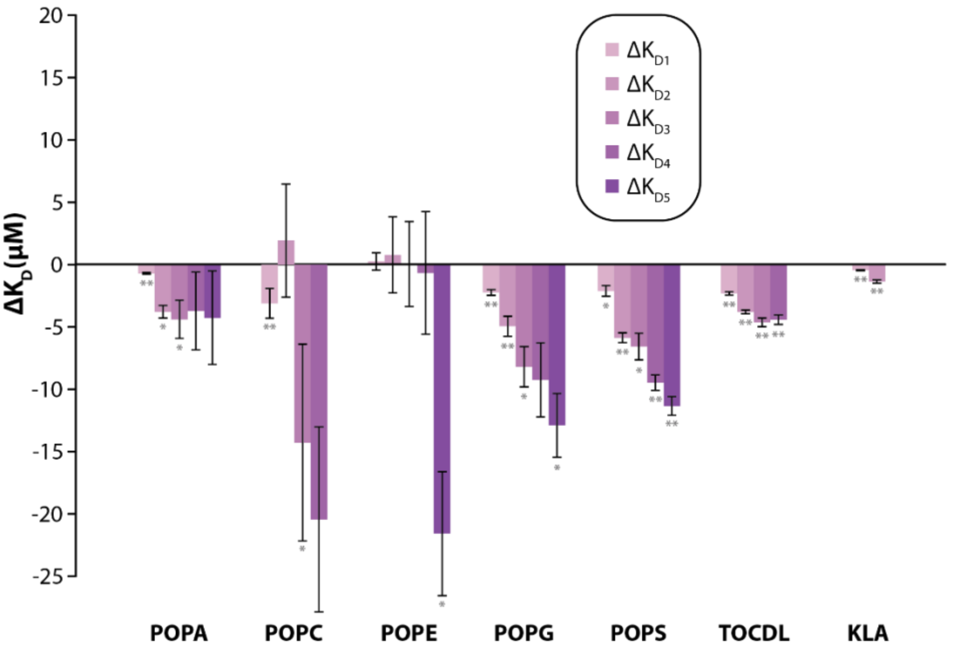


**Figure S14. The conformation of MsbA influences lipid binding affinity.** Reported are the difference in the trapped minus non-trapped $K_D$ values. The asteriks denote student's t-test for MsbA partially versus fully loaded with copper(II) (*$p$<0.05, **$p$<0.01).

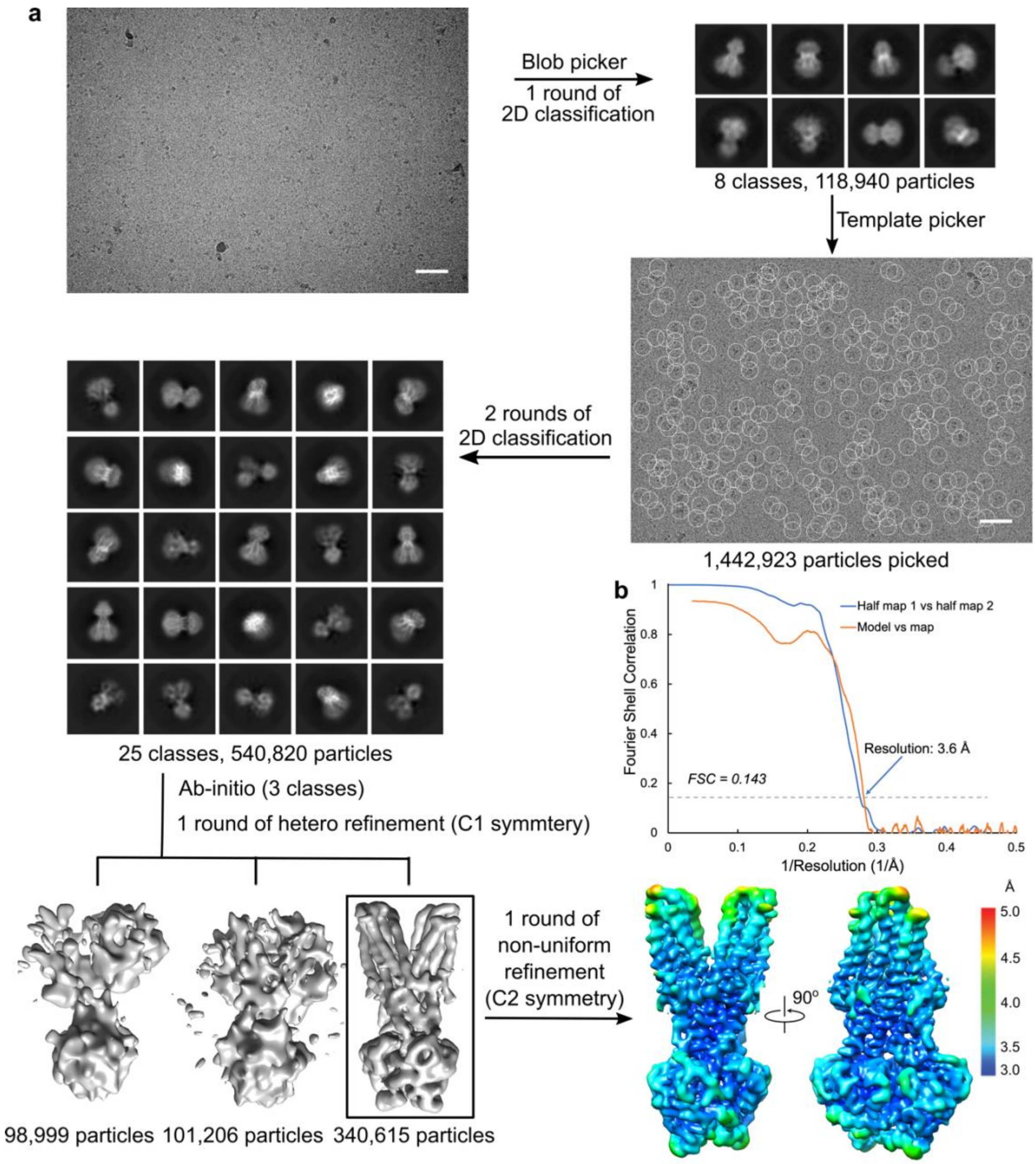


**Figure S15. Single-particle cryoEM anlaysis for MsbA trapped with vanadate and in complex with KLA.** A) The workflow of data processing. Initial motion correction was carried out using MotionCor2.[9] A representative motion-corrected micrograph is shown along with a 50-nm scale bar. The following automatic picking and 2D classification were performed in cryoSPARC.[10] Particle selection was performed using the 2D templates generated by blob picker, followed by a 2D classification. Representative 2D class averages are shown, with the box edge corresponding to 213 Å. After disposing contamination and poorly-aligned 2D classes, the particles were classified by hetero refinement with 3 initial models generated using ab initio reconstructions. A non-uniform refinement was performed on the best class of particles with a C2 symmetry. B) Fourier shell correlation curves of the final reconstruction and model versus map. The resolution of the reconstruction was determined by the FSC=0.143 criterion.

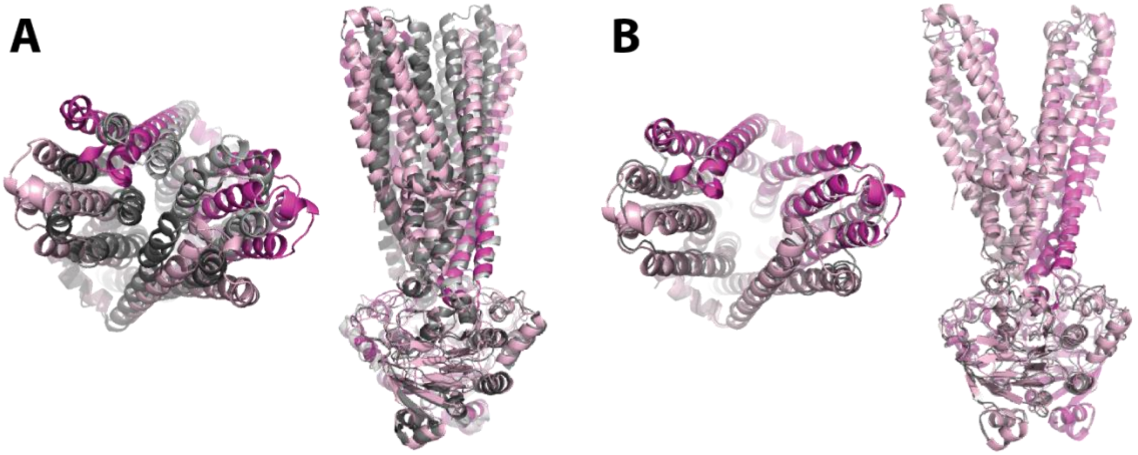


**Figure S16. Comparison of the vanadate-trapped MsbA structure to other structures.** A) Alignment of outward, occluded MsbA structure (PDB 5TTP), shown in pink and magenta, with trapped structure shown in grey. B) Overlay of the MsbA structure with the open, outward facing structure of *S. typhimurium* MsbA bound to AMPPNP, a non-hydrolyzable ATP analog (PDB 3B60). Shown as described for panel A.

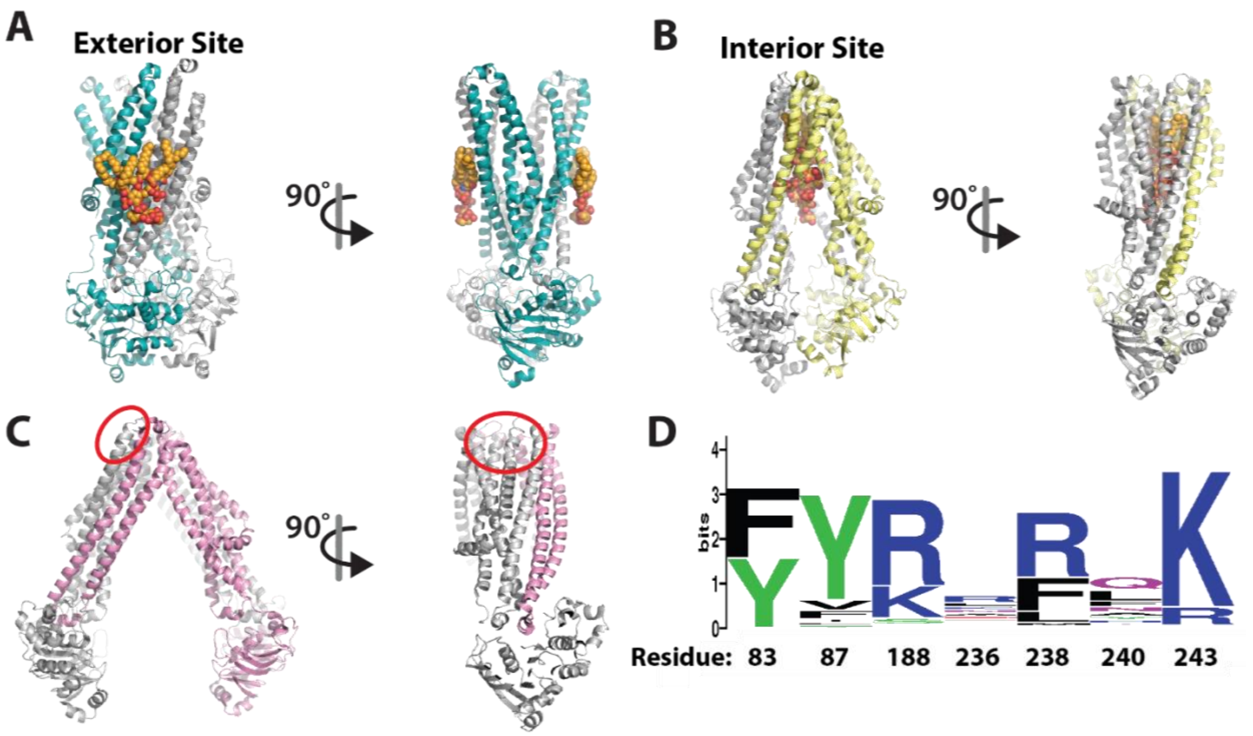


**Figure S17. Comparison of reported and conservation of MsbA LPS binding sites**. Structures are shown in cartoon representation with lipid (if present) shown in orange spheres. Two views are shown (0 and 90°). A-C) Shown is the A) exterior binding site (this work), B) interior binding site (PDB 6BPL),[11,20] and C) putative exterior binding site (PDB 6BL6).[21,22] In panel C, the density was not clear enough to model the lipid and putative site is denoted by a red circle.[21] D) Sequence logo of KLA interacting residues (83, 87, 188, 236, 238, 240, and 243) based on an alignment of 258 MsbA sequences from across the bacterial phylogeny.

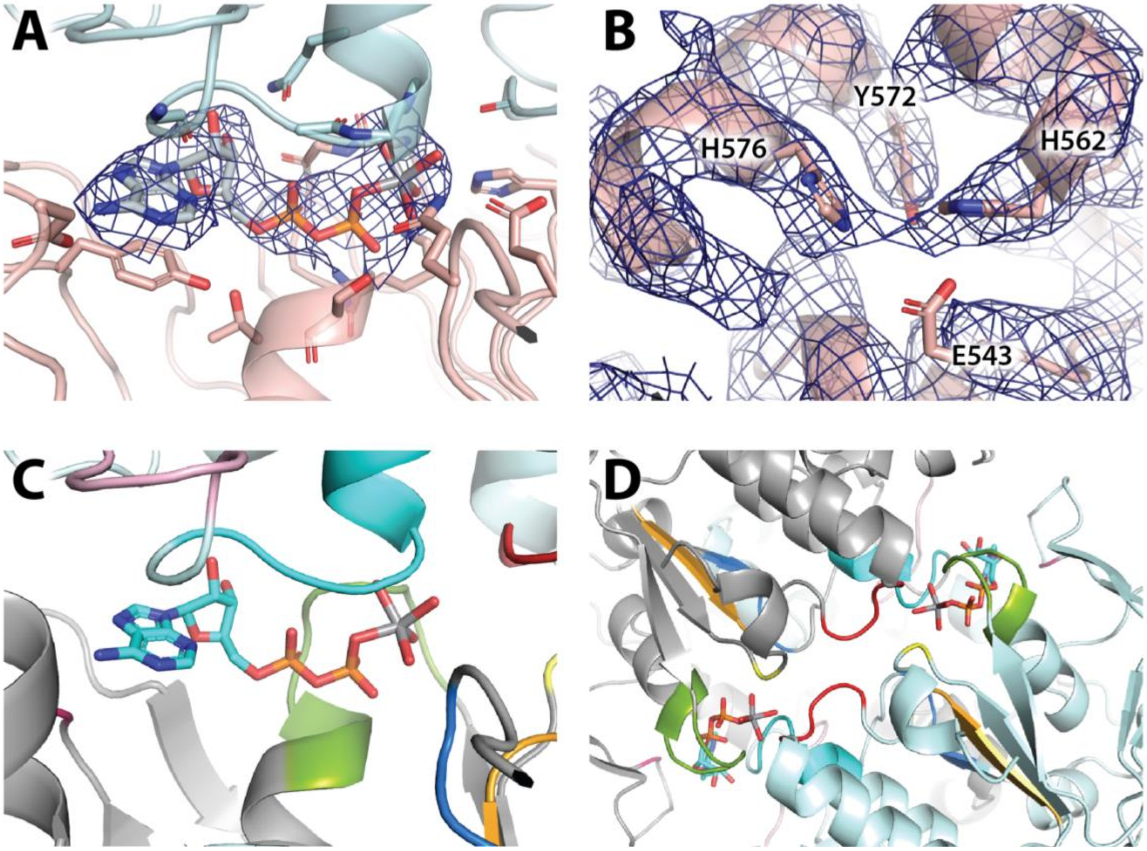


**Figure S18. CryoEM density and conserved regions of MsbA bound to ADP and vanadate.** A) The same view of the bound nucleotide as shown in Figure 4E with cryoEM density map contoured at 6 rmsd. B) A previously proposed metal binding site with density map shown as in panel A. Copper is not bound at this location. C-D) Conserved motifs in the ABC transporters. Highlighted are the hydrophobic residue of A-loop (residue 351) in pink, Walker A motif (GxxGxGK(S/T) where x denotes a hydrophobic residue, resides 376-383) or P-loop in chartreuse, Q-loop ( $\phi(\phi/Q)Q$ where denotes a hydrophobic residue, residues 421-424), X-loop (residues 472-478) in light pink, C-loop motif (consensus sequence of LSGGQ, residues 481-487) in cyan, Walker B motif ($\phi_4D$, residues 501-506) in bright orange, D-loop (consensus sequence of SALD, residues 509-512), and the H-switch histidine (residue 537) in yellow.[23-25] The residue(s) corresponding to the conserved motifs are noted.

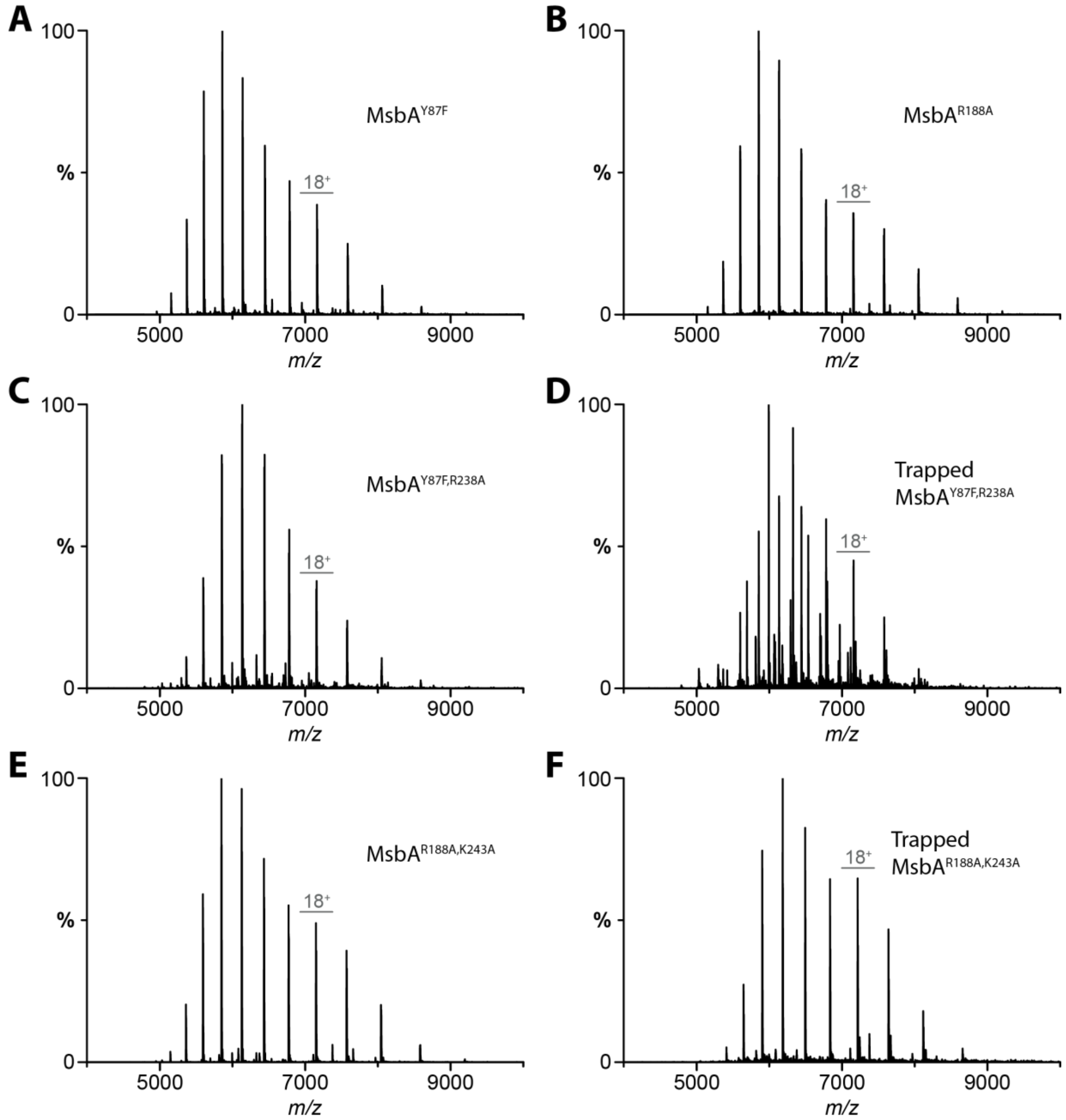


**Figure S19. Representative native mass spectra of MsbA mutants.** A) MsbA$^{Y87F}$ B) MsbA$^{R188A}$ C) MsbA$^{Y87F,R238A}$ D) MsbA$^{Y87F,R238A}$ trapped with ADP and vanadate E) MsbA$^{R188A,K243A}$ F) MsbA$^{R188A,K243A}$ trapped with ADP and vanadate.

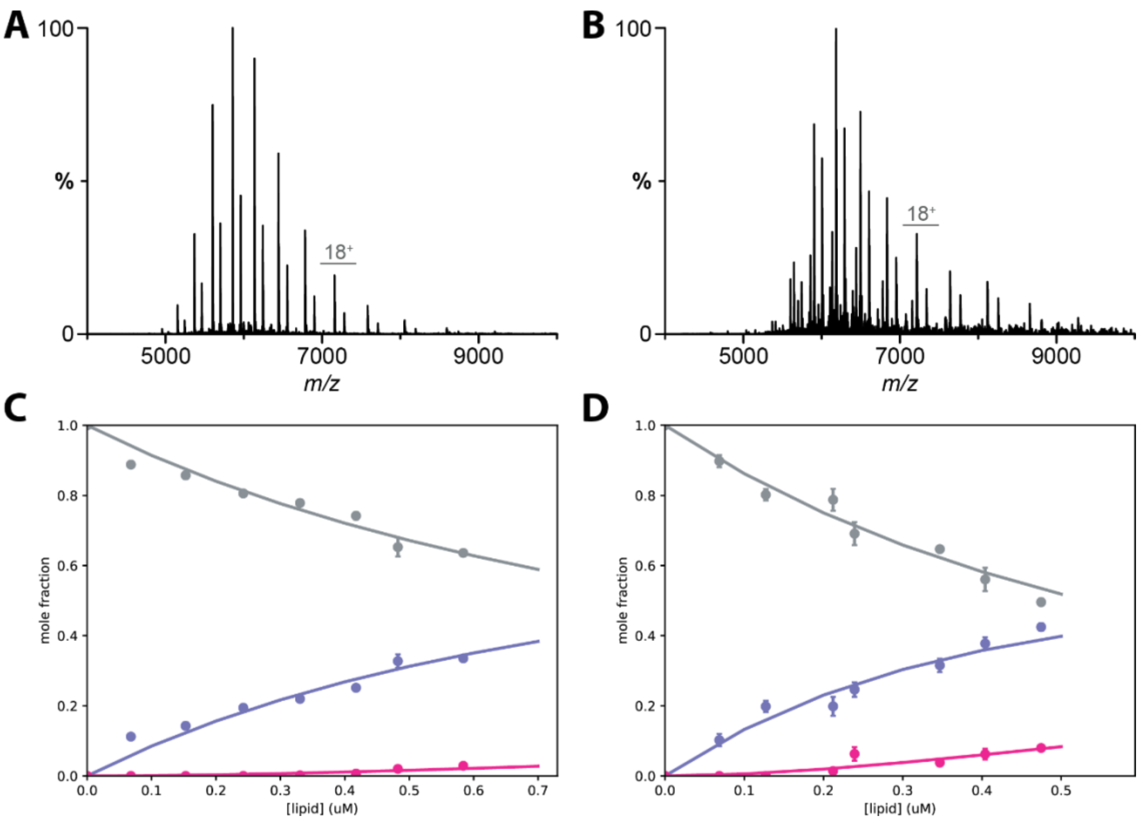


**Figure S20. Native mass spectra and plot of mole fraction for KLA binding to MsbA$^{R188A,R243A}$.** A) Native mass spectrum of 0.3 µM of MsbA$^{R188A,R243A}$ mixed with 0.7 µM of KLA. B) Native mass spectrum of 0.38 µM of MsbA$^{R188A,R243A}$ trapped with ADP and vanadate and mixed with 0.7 µM of KLA. Mole fraction plots of C) apo and D) vanadate-trapped MsbA$^{R188A,R243A}$ binding to different numbers of KLA. Shown as described in Figure S4.

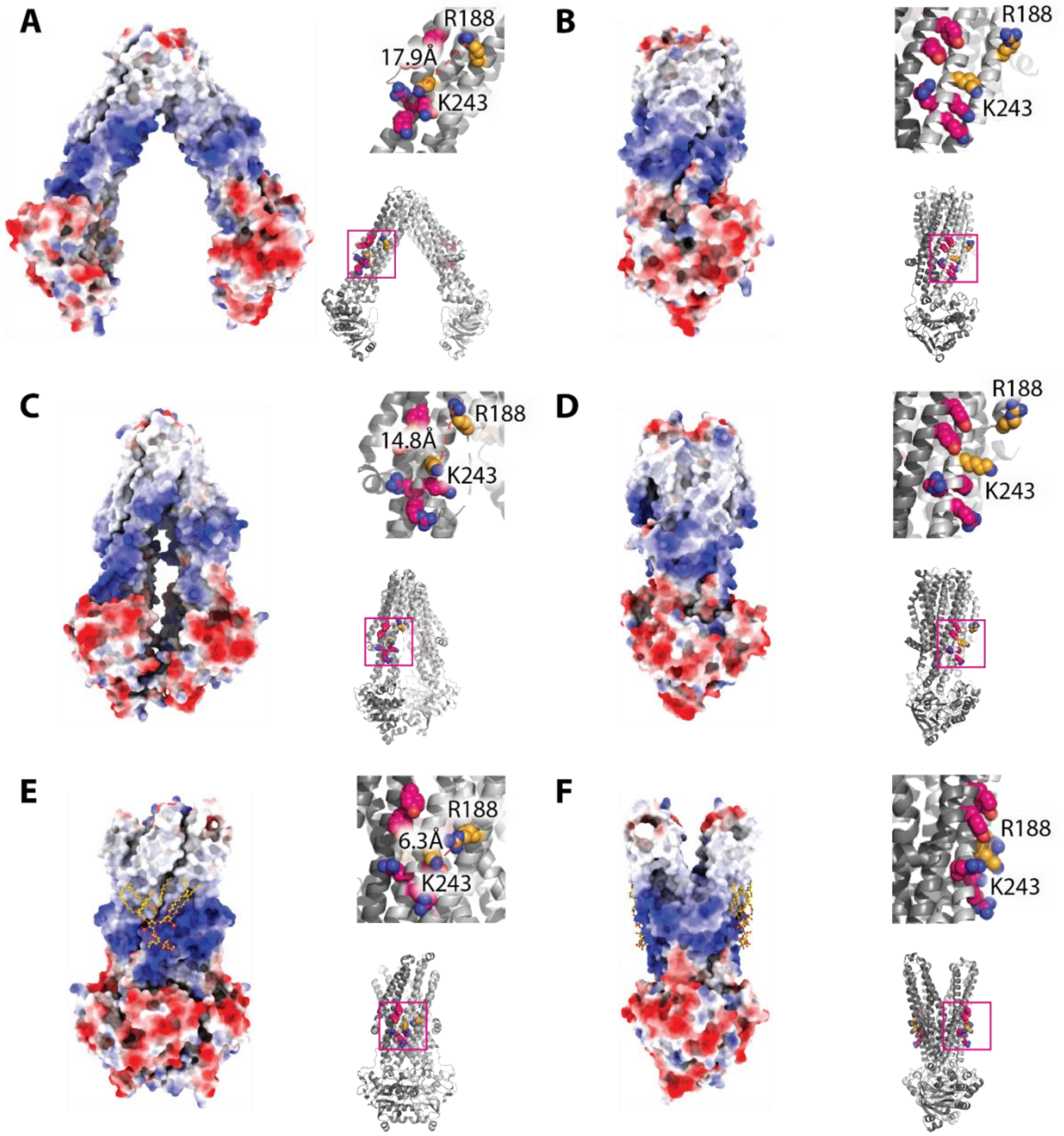


**Figure S21. Electrostatics of MsbA in different conformations.** Shown are 0 and 90° views of A-B) open, inward-facing (this work), C-D) occluded, inward-facing and bound to G907 inhibitor (PDB 6BPL), and E-F) open, outward-facing (this work) conformation. The insets show KLA binding residues in ball and stick representation, and the distance between Cζ of R188 and Nζ of K243 is labeled.

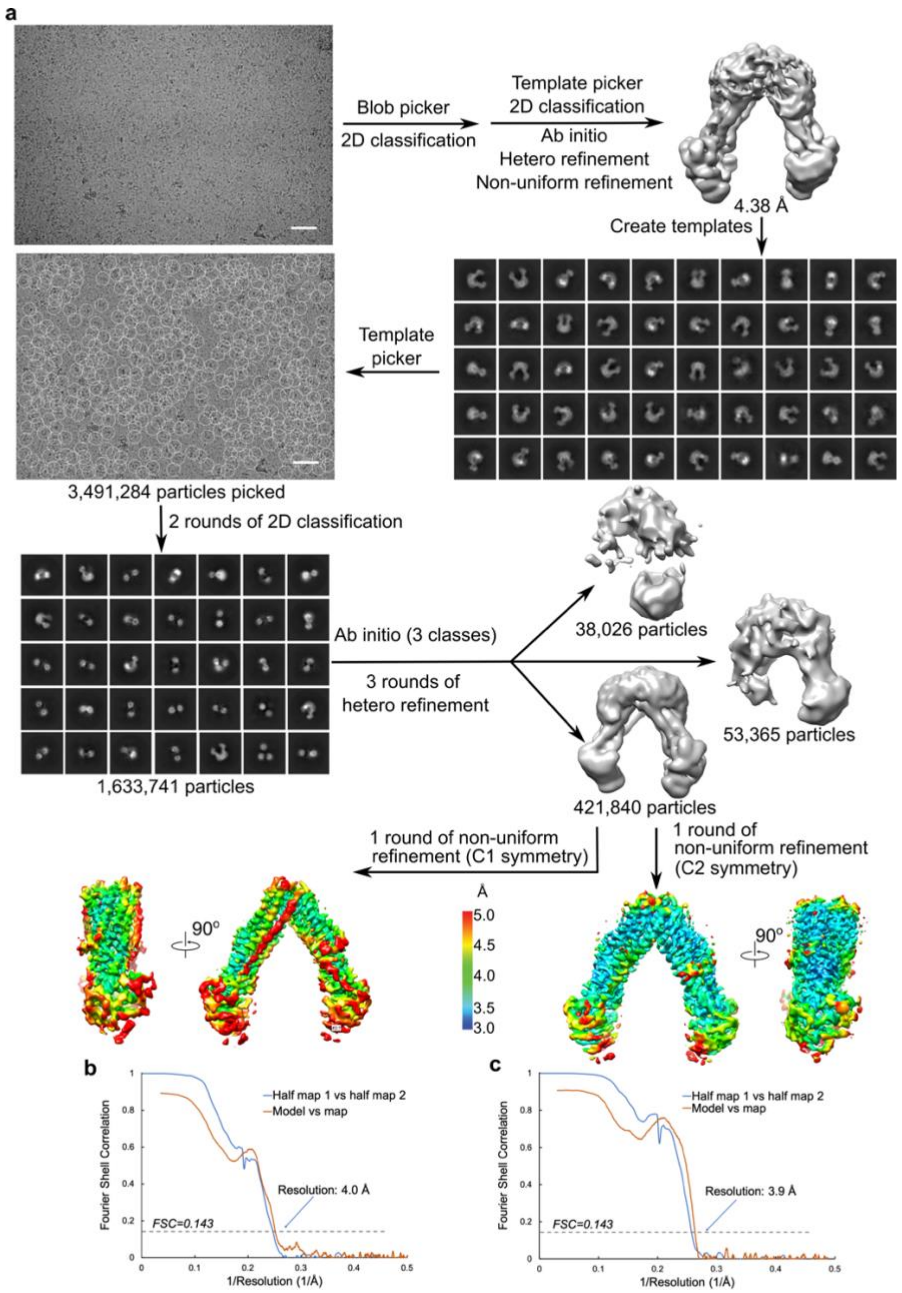


**Figure S22. Single-particle cryoEM anlaysis for open, inward-facing MsbA.** A) The workflow of data processing. The representative micrograph is shown along with a 50-nm scale bar. Particle picking was performed using the 2D templates generated by a reconstructed model from earlier runs. Representative 2D class averages are shown, with the box edge corresponding to 298 Å. After disposing contamination and poorly-aligned 2D classes, the particles were classified by hetero refinement with 3 initial models generated using ab initio reconstructions. A non-uniform refinement was performed on the best class of particles with either C1 or C2 symmetry imposed. B-C) Fourier shell correlation curves of the final reconstruction and model versus map for C1 (B) and C2 (C), respectively. The resolution of the reconstruction was determined by the FSC=0.143 criterion.

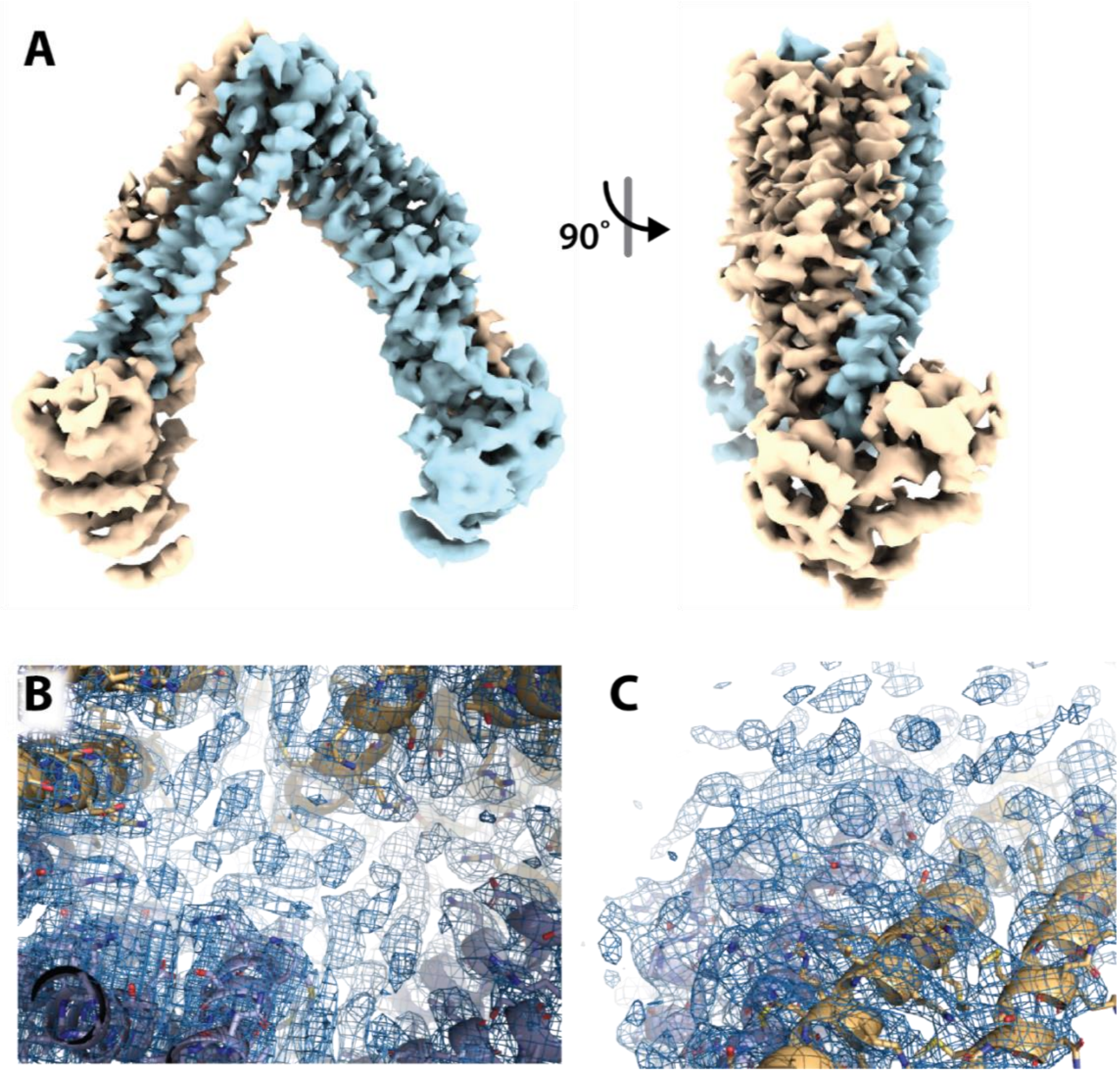


**Figure S23. CryoEM Structure of apo MsbA in an open, inward facing conformation.** A) CryoEM reconstruction (3.8 Å) with the subunits are colored beige and blue. B-C) CryoEM maps showing views B) around residue 87 and C) interior cavity with tube-like density.

### Supplementary Tables

**Table S1. Theoretical and experimental masses of MsbA.** Reported are the mean and standard deviation of centroid masses.

| | Theoretical (Da) | Experimental (Da) |
|---|---|---|
| **$MsbA_2$** | 128,947 | 128,948 ± 7 |
| **$MsbA_2(Cu)_1$** | 129,011 | 129,012 ± 5 |
| **$MsbA_2(Cu)_2$** | 129,074 | 129,079 ± 1 |
| **$MsbA_2(ADP)_2(VO_4)_2$** | 130,032 | 130,005 ± 6 |
| **$MsbA_2(ADP)_2(VO_4)_2(Cu)_1$** | 130,095 | 130,070 ± 4 |
| **$MsbA_2(ADP)_2(VO_4)_2(Cu)_2$** | 130,159 | 130,139 ± 3 |

**Table S2. Operating parameters for ICP-MS analysis of MsbA samples.**

| Parameters | |
|---|---|
| **RF Power** | 1600 W |
| **Plasma Ar Flow** | 18.0 L $min^{-1}$ |
| **Auxiliary Ar Flow** | 1.20 L $min^{-1}$ |
| **Nebulizer Ar Flow** | 0.98 L $min^{-1}$ |
| **Sample Introduction System** | Meinhard concentric nebulizer with cyclonic spray chamber |
| **Operating Frequency** | 40 MHz |
| **Sample Uptake Rate** | 1 mL $min^{-1}$ |
| **Detector Mode** | Analog |
| **Sampler/Skimmer/Hyper-skimmer Cones** | Ni/Ni/Al |
| **Scanning Mode** | Peak hopping |
| **Number of points per peak** | 5 |
| **Dwell Time** | 50 ms |
| **Sweeps per Reading** | 20 |
| **Isotopes** | $^{51}V$, $^{60}Ni$, $^{63}Cu$, $^{66}Zn$ |
| **Internal Standards** | $^{115}In$ |
| **Software** | Syngistix Software, Version 2.4 |

**Table S3. Results from ICP-MS analysis of MsbA samples.**

| Sample | V[ng/mL] | Ni[ng/mL] | Cu[ng/mL] | Zn [ng/mL] |
|---|---|---|---|---|
| **MsbA** | 38 | 10 | 1040 | 7.4 |
| **Vanadate-trapped MsbA** | 8338 | 16 | 1313 | 11 |

**Table S4. Equilibrium binding constants for lipids binding to MsbA partially loaded with copper(II).** $K_D$ was obtained from fitting a sequential lipid binding model to the mole fraction data. Reported are the mean and standard deviation (*n* = 3).

| | $K_{D1}$(μM) | $K_{D2}$(μM) | $K_{D3}$(μM) | $K_{D4}$(μM) | $K_{D5}$(μM) | $K_{D6}$(μM) | $K_{D7}$(μM) | $K_{D8}$(μM) | $K_{D9}$(μM) | $R^{2*}$ | $X^{2*}$ |
|---|---|---|---|---|---|---|---|---|---|---|---|
| **POPA** | 14.7 ± 1.5 | 23.2 ± 0.6 | 28.5 ± 2.7 | 30.8 ± 2.7 | 32.6 ± 1.9 | | | | | 0.98 | 0.07 |
| **POPC** | 20.4 ± 1.4 | 39.5 ± 2.0 | 46.0 ± 4.6 | 47.0 ± 9.4 | 34.2 ± 4.6 | | | | | 0.98 | 0.08 |
| **POPE** | 12.4 ± 0.7 | 23.9 ± 2.4 | 31.5 ± 3.5 | 32.5 ± 6.5 | 33.7 ± 6.9 | 32.9 ± 4.9 | 28.0 ± 5.0 | | | 0.98 | 0.08 |
| **POPG** | 4.9 ± 0.4 | 11.1 ± 0.8 | 17.4 ± 1.7 | 23.1 ± 0.6 | 27.1 ± 0.9 | 28.6 ± 2.0 | 32.5 ± 3.0 | | | 0.95 | 0.11 |
| **POPS** | 3.6 ± 0.2 | 8.8 ± 0.05 | 14.7 ± 0.5 | 20.2 ± 1.3 | 22.2 ± 1.1 | 23.1 ± 1.1 | 25.1 ± 1.0 | 34.1 ± 1.9 | 36.7 ± 6.7 | 0.96 | 0.10 |
| **TOCDL** | 1.9 ± 0.1 | 4.3 ± 0.7 | 6.7 ± 0.9 | 9.4 ± 1.5 | 12.5 ± 4.0 | 9.4 ± 1.6 | | | | 0.99 | 0.04 |
| **KLA** | 0.9 ± 0.1 | 2.0 ± 0.4 | 3.5 ± 0.4 | 5.0 ± 0.3 | | | | | | 0.91 | 0.26 |

*These values represent the replicates with the poorest fits.

**Table S5. Equilibrium binding constants for lipids binding to MsbA saturated with copper(II).** Reported as described in Table S4.

| | $K_{D1}$(μM) | $K_{D2}$(μM) | $K_{D3}$(μM) | $K_{D4}$(μM) | $K_{D5}$(μM) | $K_{D6}$(μM) | $K_{D7}$(μM) | $K_{D8}$(μM) | $K_{D9}$(μM) | $R^{2*}$ | $X^{2*}$ |
|---|---|---|---|---|---|---|---|---|---|---|---|
| **POPA** | 2.6 ± 0.04 | 5.7 ± 0.1 | 8.5 ± 0.3 | 10.5 ± 0.4 | 14.0 ± 0.5 | 16.9 ± 0.3 | 23.4 ± 3.8 | | | 0.99 | 0.02 |
| **POPC** | 17.2 ± 0.6 | 33.6 ± 1.3 | 48.5 ± 4.1 | 53.7 ± 4.9 | | | | | | 0.99 | 0.02 |
| **POPE** | 7.4 ± 0.3 | 18.4 ± 0.7 | 27.8 ± 0.5 | 33.1 ± 2.0 | 39.8 ± 3.2 | 37.8 ± 4.0 | | | | 0.99 | 0.02 |
| **POPG** | 4.1 ± 0.1 | 10.5 ± 0.2 | 16.2 ± 0.3 | 20.6 ± 0.7 | 25.4 ± 1.1 | 27.6 ± 1.8 | 31.7 ± 1.6 | 36.4 ± 1.2 | 36.4 ± 2.7 | 1.00 | 0.01 |
| **POPS** | 3.7 ± 0.04 | 5.7 ± 0.2 | 9.3 ± 0.6 | 13.0 ± 0.4 | 16.8 ± 0.5 | 19.8 ± 0.5 | 22.0 ± 0.6 | 25.4 ± 0.7 | 27.8 ± 0.1 | 0.99 | 0.03 |
| **TOCDL** | 1.6 ± 0.08 | 2.9 ± 0.09 | 4.1 ± 0.2 | 5.1 ± 0.2 | | | | | | 1.00 | 0.01 |
| **KLA** | 0.6 ± 0.02 | 2.0 ± 0.09 | | | | | | | | 0.98 | 0.04 |

*These values represent the replicates with the poorest fits.

**Table S6. Summary of X-ray data collection and refinement statistics.**

| N-term MsbA fusion to GFP in complex with copper(II) | | |
|---|---|---|
| **Crystal Parameters** | | |
| **Space Group** | $I2_12_12_1$ | $I2_12_12_1$ |
| **Unit Cell Dimensions** | *a*=70.61, *b*=108.54, *c*=140.4 | *a*=70.62, *b*=108.96, *c*=140.64 |
| **Angles** | α=β=γ=90° | α=β=γ=90° |
| **Data Collection** | | |
| **Synchrotron (Beamline)** | APS (24-ID-C) | TAMU (R-Axis IV++) |
| **λ (Å)** | 1.378 | 1.54 |
| **Resolution (Å)** | 42.98- 2.15 (2.21- 2.15) | 70.32-2.2 (2.28-2.2) |
| **$I/\sigma$** | 12.35 (1.61) | 12.17 (2.14) |
| **Observed Reflections** | 60359 (4433) | 376860 (27999) |
| **Unique Reflections** | 17601 (1280) | 56321 (4109) |
| **Completeness (%)** | 99.3 (98.0) | 99.2 (98.6) |
| **Refinement** | | |
| **$R_{work}$ (%)** | 0.19 (0.29) | 0.19 (0.34) |
| **$R_{free}$ (%)** | 0.21 (0.30) | 0.22 (0.34) |
| **Number of Protein Atoms** | 1968 | 1955 |
| **Nonprotein Atoms** | 26 | 23 |
| **Number of water molecules** | 48 | 52 |
| **Number of protein residues** | 236 | 236 |
| **RMS (bonds)** | 0.008 | 0.007 |
| **RMS (angles)** | 0.91 | 0.88 |
| **Ramachandran favored (%)** | 96.98 | 97.41 |
| **Ramachandran allowed (%)** | 3.02 | 2.59 |
| **Ramachandran outliers (%)** | 0.00 | 0.00 |
| **Rotamer outliers (%)** | 0.96 | 2.43 |
| **Wilson B-factor** | 52.33 | 57.39 |
| **B factor of protein atoms** | 62.70 | 63.71 |
| **PDB Code** | 8DHY | |

**Table S7. Equilibrium binding constants for lipids binding to MsbA trapped with ADP and vanadate.** Reported as described in Table S4.

| | $K_{D1}$(µM) | $K_{D2}$(µM) | $K_{D3}$(µM) | $K_{D4}$(µM) | $K_{D5}$(µM) | $K_{D6}$(µM) | $K_{D7}$(µM) | $K_{D8}$(µM) | $K_{D9}$(µM) | $R^{2*}$ | $X^{2*}$ |
|---|---|---|---|---|---|---|---|---|---|---|---|
| **POPA** | 2.2 ± 0.01 | 3.2 ± 0.3 | 5.6 ± 10.0 | 8.0 ± 2.0 | 11.2 ± 2.4 | 14.3 ± 2.3 | | | | 0.84 | 0.28 |
| **POPC** | 15.1 ± 0.5 | 34.9 ± 2.7 | 39.0 ± 3.3 | 40.2 ± 0.3 | 33.2 ± 3.8 | | | | | 0.95 | 0.15 |
| **POPE** | 7.5 ± 0.4 | 18.9 ± 1.9 | 27.8 ± 2.2 | 32.7 ± 2.6 | 25.4 ± 0.9 | 25.2 ± 5.7 | | | | 0.95 | 0.13 |
| **POPG** | 2.6 ± 0.1 | 7.2 ± 0.5 | 10.8 ± 1.0 | 14.5 ± 1.8 | 16.8 ± 1.3 | 21.4 ± 0.5 | 21.1 ± 0.7 | 23.4 ± 1.5 | 27.0 ± 0.6 | 0.93 | 0.14 |
| **POPS** | 2.3 ± 0.3 | 1.8 ± 0.2 | 4.9 ± 0.4 | 6.7 ± 0.06 | 9.3 ± 0.2 | 13.5 ± 0.2 | 16.2 ± 0.7 | 20.9 ± 0.6 | 20.3 ± 1.6 | 0.93 | 0.14 |
| **TOCDL** | 0.1 ± 0.04 | 0.4 ± 0.04 | 1.0 ± 0.04 | 2.1 ± 0.1 | | | | | | 0.86 | 0.33 |
| **KLA** | 0.3 ± 0.02 | 1.1 ± 0.01 | | | | | | | | 0.97 | 0.06 |

*These values represent the replicates with the poorest fits.

**Table S8.** **Statistics of cryo-EM data collection and processing.**

| | MsbA trapped with ADP-vanadate and bound to KLA | Open, inward-facing MsbA |
|---|---|---|
| **Microscope** | Krios (University of Chicago) | Krios (University of Chicago) |
| **Magnification** | 81,000 | 81,000 |
| **Voltage (kV)** | 300 | 300 |
| **Spherical aberration (mm)** | 2.7 | 2.7 |
| **Detector** | K3 | K3 |
| **Camera mode** | Super resolution counting | Super resolution counting |
| **Exposure rate ($e^-$/pixel/s)** | 15 | 15 |
| **Total exposure ($e^-$/Å$^2$)** | 50 | 50 |
| **Defocus range (μm)** | -1.0 to -2.5 | -1.0 to -2.5 |
| **Pixel size (Å)** | 0.5325 (1.065 physical) | 0.5325 (1.065 physical) |
| **Mode of data collection** | Image shift | Image shift |
| **Energy filter** | 20 eV slit | 20 eV slit |
| **Software for data collection** | EPU | EPU |
| **Number of micrographs** | 4,221 | 5,530 |
| **Symmetry imposed** | C2 | C2 |
| **Box size (pixel)** | 200 | 280 |
| **Initial particle images (no.)** | 1,442,923 | 3,491,284 |
| **Particle images for 3D (no.)** | 540,820 | 1,633,741 |
| **Final particle images (no.)** | 340,615 | 421,840 |
| **Map resolution, unmasked (Å)** | 4.0 | 4.2 |
| **Map resolution, masked (Å)** | 3.6 | 3.9 |
| **B-factor used for sharpening (Å$^2$)** | 213.7 | 208.6 |
| **EMD accession code** | EMD-27544 | EMD-27545 |

**Table S9. Statistics of cryo-EM model refinement and geometry for vanadate-trapped MsbA**

| Model | MsbA trapped with ADP-vanadate and bound to KLA | open, inward facing MsbA |
|---|---|---|
| **Composition (#)** | | |
| **Chains** | 2 | 2 |
| **Atoms** | 9314 (Hydrogens: 0) | 8912 (Hydrogens: 0) |
| **Residues** | Protein: 1152 Nucleotide: 0 | Protein: 1150 Nucleotide: 0 |
| **Water** | 0 | 0 |
| **Ligands** | KLA: 2 | 0 |
| | AOV: 2 | |
| **Bonds (RMSD)** | | |
| **Length (Å) (# > 4σ)** | 0.004 (0) | 0.003 (0) |
| **Angles (°) (# > 4σ)** | 0.997 (2) | 0.626 (0) |
| **MolProbity score** | 1.52 | 1.66 |
| **Clash score** | 9.94 | 8.91 |
| **Ramachandran plot (%)** | | |
| **Outliers** | 0.00 | 0.09 |
| **Allowed** | 1.13 | 2.97 |
| **Favored** | 98.87 | 96.95 |
| **Rama-Z (Ramachandran plot Z-score, RMSD)** | | |
| **whole** | (N = 1148) 1.55 (0.26) | (N = 1146) 1.76 (0.25) |
| **helix** | (N = 728) 1.68 (0.20) | (N = 700) 2.03 (0.20) |
| **sheet** | (N = 50) -0.04 (0.72) | (N = 74) 1.25 (0.61) |
| **loop** | (N = 370) -0.03 (0.35) | (N = 372) -0.50 (0.32) |
| **Rotamer outliers (%)** | 0.00 | 0.00 |
| **Cβ outliers (%)** | 0.00 | 0.00 |
| **Peptide plane (%)** | | |
| **Cis proline/general** | 0.0/0.0 | 0.0/0.0 |
| **Twisted proline/general** | 0.0/0.0 | 0.0/0.0 |
| **CaBLAM outliers (%)** | 0.52 | 1.58 |
| **ADP (B-factors)** | | |
| **Iso/Aniso (#)** | 9314/0 | 8912/0 |
| **min/max/mean** | | |
| **Protein** | 21.49/152.72/67.77 | 0.17/104.70/43.85 |
| **Ligand** | 53.29/79.81/72.96 | |
| **Data** | | |
| **Box** | | |
| **Lengths (Å)** | 75.62, 86.27, 137.39 | 89.46, 136.32, 136.32 |
| **Angles (°)** | 90.00, 90.00, 90.00 | 90.00, 90.00, 90.00 |
| **Supplied Resolution (Å)** | 3.6 | 3.9 |
| **Resolution Estimates (Å)** | Masked | Masked |
| **d FSC (half maps; 0.143)** | 3.6 | 3.9 |
| **d 99 (full/half1/half2)** | 3.8/4.5/4.6 | 4.1/4.6/4.7 |
| **d model** | 3.8 | 4.0 |
| **d FSC model (0/0.143/0.5)** | 3.5/3.6/3.8 | 3.7/3.8/4.0 |
| **Map min/max/mean** | -1.00/1.42/0.04 | -1.28/1.91/0.02 |
| **Model vs. Data** | | |
| **CC (mask)** | 0.81 | 0.77 |
| **CC (box)** | 0.71 | 0.73 |
| **CC (peaks)** | 0.67 | 0.66 |
| **CC (volume)** | 0.78 | 0.74 |
| **Mean CC for ligands** | 0.72 | - |
